# *In vivo* evidence that *SORL1*, encoding the endosomal recycling receptor SORLA, can function as a causal gene in Alzheimer’s Disease

**DOI:** 10.1101/2021.07.13.452149

**Authors:** Olav M. Andersen, Nikolaj Bøgh, Anne M. Landau, Gro Grunnet Pløen, Anne Mette G. Jensen, Giulia Monti, Benedicte Parm Ulhøi, Jens Randel Nyengaard, Kirsten Rosenmay Jacobsen, Margarita Melnikova Jørgensen, Ida E. Holm, Marianne L. Kristensen, Esben Søvsø Szocska Hansen, Charlotte E. Teunissen, Laura Breidenbach, Mathias Droescher, Ying Liu, Hanne Skovsgaard Pedersen, Henrik Callesen, Yonglun Luo, Lars Bolund, David J. Brooks, Christoffer Laustsen, Scott A. Small, Lars F. Mikkelsen, Charlotte B. Sørensen

## Abstract

The few established causal genes in Alzheimer’s disease (AD), mutations in *APP* and *PSENs,* have been functionally characterized using biomarkers, capturing an *in vivo* profile reflecting the disease’s initial preclinical phase. *SORL1*, a gene encoding the endosome recycling receptor SORLA, epidemiologically behaves as a causal gene when truncating mutations lead to partial loss of protein function. Here, in an effort to test whether *SORL1* can indeed function as an AD causal gene, we used CRISPR-Cas9-based gene editing to develop a novel model of *SORL1* haploinsufficiency in Göttingen Minipigs taking advantage of porcine models for biomarker investigations. *SORL1* haploinsufficiency in young minipigs was found to phenocopy the preclinical *in vivo* profile of AD observed with other causal genes, resulting in spinal fluid abnormalities in Aβ and tau, with no evident neurodegeneration or amyloid plaque formation. These studies provide functional support that *SORL1* is a bona fide causal gene in AD, and when taken together with recent insight on other AD-causal genes, support the idea that dysfunctional endosomal recycling is a dominant pathogenic pathway in the disease.

## INTRODUCTION

Whereas old age is among the strongest risk factors for development of Alzheimer’s disease (AD), there is also evidence of genetic components of the disease. Causal genes in AD turn out, however, to be extremely rare, with most AD-associated genes affecting disease risk. “Risk” is on a continuum and, epidemiologically, only those genetic mutations which at population levels show extremely high effects sizes and odds ratios can be considered causal in AD pathogenesis. Mechanistically, causal genes are those who by themselves can trigger the disease’s initiating pathologies, while risk genes do not necessarily act as pathogenic triggers, but more typically modulate how the brain responds to them.

Until recently, only three genes harbor specific mutations that fulfill both epidemiological and mechanistic criteria, justifying their designation as causal genes: *APP* (encoding the amyloid precursor protein), *PSEN1* (presenilin1) and *PSEN2* (presenilin 2) (1). Recently *SORL1* (sortilin related receptor 1) has been proposed to be included to the exclusive list of AD causal genes (2). Patients harboring mutations in *APP* or *PSEN1/2* have been investigated by cerebrospinal fluid (CSF) analysis and neuroimaging, thereby establishing an *in vivo* signature of the earliest preclinical stage of disease characterized by CSF elevations of APP’s amyloid β-peptide (Aβ) and tau, before the onset of amyloid plaques and neurodegeneration (3, 4). This initial and presumably initiating pathogenic profile was confirmed in mouse models genetically engineered to express these mutations (5), an impotent validation because of the complexity of tracking the onset and progression of AD’s slowly worsening pathophysiology via CSF analyses and imaging in mice.

Endosome dysfunction is strongly implicated in the underlying pathogenesis of both sporadic and familiar forms of AD (6, 7). Relying on studies suggesting that the *SORL1*-encoded protein SORLA can act as a receptor of the endosome’s retromer complex (8, 9), the first genetic linkage of *SORL1* to AD emerged from an association analysis that investigated genes encoding components of the retromer complex, *SORL1*, and other retromer-related receptors (10). Since then, numerous studies have confirmed *SORL1*’s genetic link to AD through hypothesis-free genome and exome screening (11, 12). Most recently, genetic epidemiology studies have identified rare truncation mutations in *SORL1* almost exclusively in AD patients suggesting that *SORL1* functions as a causal gene (12).

In parallel, numerous subsequent studies have further confirmed that SORLA functions in endosomal recycling of cargo proteins (13–15), which when disrupted is a dominant upstream pathogenic pathway in AD (16).

Because *SORL1* truncation mutations typically cause partial loss of protein function, inducing protein haploinsufficiency has turned out to be a valid and convenient model of these mutations. Indeed, previous work using mouse models or neuronal culture studies have shown that *SORL1* deficiency can phenocopy the cell biology of the other causal genes (i.e. *APP*, *PSEN1* and *PSEN2*). In particular, and just like the disease-causing variants of these causal genes, *SORL1* deficiency has been shown to accelerate Aβ production (17, 18) as well as causing abnormal swelling of neuronal endosomes (19), both hallmark features of sporadic and familiar forms of AD (20).

Since these studies were not, and cannot, be performed in patients, it remains unknown whether *SORL1* loss-of-function phenocopies the *in vivo* preclinical signature (i.e. increased CSF levels of Aβ and tau before the onset of amyloid plaques and neurodegeneration) of the established causal genes. Multiple large-scale and long-term global consortia were required to establish this earliest preclinical *in vivo* signature of AD in patients harboring the established causal genes. It will take an equally extensive infrastructure and long time-course to ultimately test whether patients harboring the putatively causal *SORL1* genetic variants phenocopy the *in vivo* preclinical signature. Until then, as with *APP* and the *PSEN*s, this signature can be tested for in appropriate animal models.

Here we investigated the causality of *SORL1* in relation to AD in a novel Göttingen Minipigs model genetically engineered to be *SORL1* haploinsufficient due to heterozygous knockout of the endogenous porcine *SORL1* gene, thus mimicking the genetic status of AD patients with heterozygous *SORL1* truncating mutations. Besides the general public acceptance of using pigs rather than non-human primates for biomedical research, pigs have, in contrast to mice, gyrencephalic brains closely resembling the human brain (21) and also share more ultra-conserved genomic regions with humans than mice (22). Importantly, the pig has a body size and longevity enabling both the ability to deploy CSF and neuroimaging markers employed in the clinical setting of AD diagnosis and allowing for longitudinal monitoring of these.

## RESULTS

### *SORL1* expression in porcine neurons

SORLA is predominantly expressed in neurons in both mouse and human brain (23). To confirm that this is also true for the pig, we examined sections of frontal cortex obtained from wild-type Göttingen Minipigs by immunohistochemistry. As expected, SORLA immunoreactivity was observed predominantly in neuronal soma (**Fig. 1a**).

**Figure 1.**
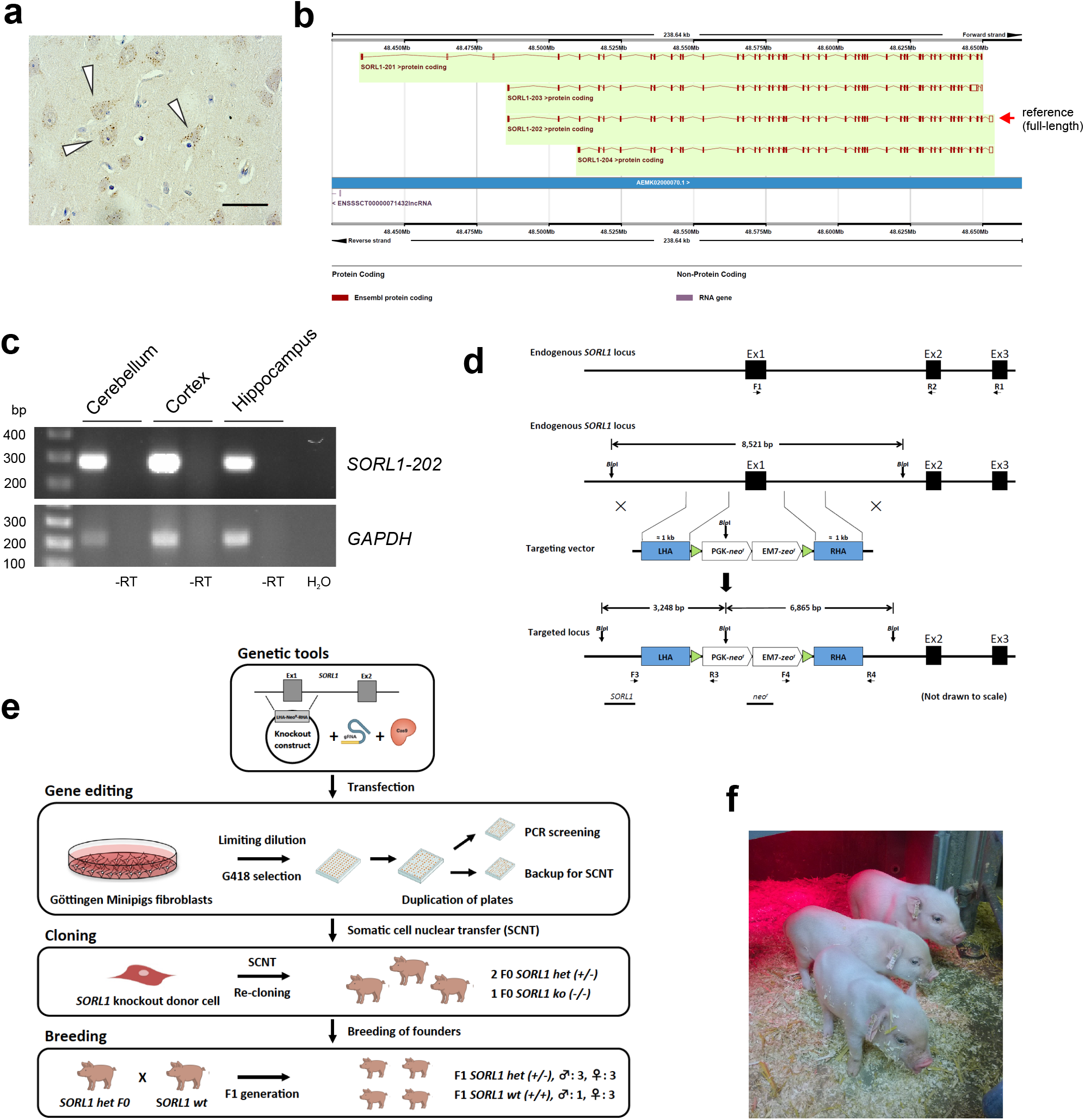
Generation of *SORL1*-deficient Göttingen Minipigs. **a)** Immunohistochemical detection of endogenous SORLA predominantly in neuronal cell soma (indicated by arrowheads) in frontal cortex from an adult *wild type* Göttingen Minipig. Scale bar is 20 μm. **b)** Schematic view of porcine *SORL1* transcripts from Ensembl Sscrofa 11.1. Four *SORL1* isoforms are listed of which *SORL1*-202, comprising 48 exons, is considered the reference transcript. **c)** RT-PCR validation of the porcine reference *SORL1*-202 transcript. cDNA obtained from reverse transcription of total RNA, isolated from cortex, hippocampus and cerebellum from a *wild type* Göttingen Minipig, was used as template in RT-PCR using primers specific for the 5’ of the reference *SORL1*-202 transcript (exon 1-3) or for porcine GAPDH. Reactions carried out in parallel using either RNA without reverse transcription (-RT) or with water (H_2_O) as templates served as negative controls. **d)** Schematic representation of the endogenous *SORL1* locus (upper panel). The three first exons of the endogenous porcine *SORL1* gene are shown as black boxes. Primers used for RT-PCR (F1+R1 and F1+R2, respectively) to validate the presence of the reference *SORL1-202* transcript in the Göttingen Minipig breed, and in *SORL1-wt*, *het* and *ko* animals, are illustrated as horizontal arrows. Schematic representation of the endogenous *SORL1* locus, the gene targeting vector and the targeted *SORL1* locus (lowest panel). The targeting vector comprises a left and right homology arm (shown as blue boxes; LHA and RHA, respectively) flanked by loxP sites (light blue triangles). The LHA and RHA are separated by a PGK-*neo^r^*/EM7-*zeo^r^* expression cassette for mammalian and bacterial selection, respectively. A region of 609 bp, comprising the entire *SORL1* exon 1 and its flanking regions, is replaced by the PGK-*neo^r^*/EM7-*zeo^r^* cassette upon successful gene editing resulting in DNA-fragments of 3,248 bp and 6,865 bp when *Blp*I-digested genomic DNA is hybridized with the *SORL1* and *neo^r^* probe, respectively. Positions of the PCR screening primer pairs F3/R3 and F4/R4 are illustrated as horizontal arrows. *Blp*I restriction sites are indicated as vertical arrows and Southern blot probes (*SORL1* ^probe^ and *neo^r^* ^probe^) are shown as black horizontal bars. **e)** Generation of *SORL1*-deficient Göttingen Minipigs. Gene editing was performed by co-transfecting primary porcine fibroblasts, isolated from newborn female Göttingen Minipigs, with the gene targeting vector, the hCas9 plasmid and the gRNA vector. Two days post transfection, the transfected cells were trypsinized and half of the cell suspension was subjected to limiting dilution by reseeding the cells into 96-well plates. Following selection for two weeks, G418-resistant cell clones were trypsinized and 1/3 of the resulting cell suspension was transferred to 96-well PCR plates for PCR screening, 1/3 was cultured further for Southern blot analysis, and 1/3 was cultured in 96-well plates for freezing at early passages and subsequent usage as nuclear donor cells for cloning of minipigs by SCNT. Two surviving piglets with heterozygous KO of *SORL1* (*SORL1 het*) and one surviving piglet with homozygous KO of *SORL1* (*SORL1 ko*) were obtained following SCNT and one re-cloning. The cloned *SORL1 het* founders (F0 generation) were used for conventional breeding to obtain *SORL1 het* and *SORL1 wt* offspring (F1 generation). **f)** Eight days old cloned Göttingen Minipigs. All piglets were born with no apparent gross abnormalities but apart from one *SORL1-ko* piglet, all *SORL1-het* piglets, died neonatally. Fibroblasts isolated from an ear biopsy from the *SORL1-het* piglet shown in front on the photo were used for re-cloning to obtain viable *SORL1-het* pigs.

The human *SORL1* gene gives rise to several protein-coding transcripts. In the pig, four *SORL1* transcripts (*SORL1*-201, 202, 203, and 204) exist according to the *Sus scrofa* genome assembly 11.1 (**Fig. 1b**). To experimentally confirm the presence of exon 1 in the reference transcript (*SORL1*-202) for subsequent gene targeting, RT-PCR analysis was performed on cDNA obtained from cortex, hippocampus and cerebellum tissues in wild-type Göttingen Minipigs using primers specific for the 5’- and 3’-end of this *SORL1* transcript. Bands corresponding to the expected size of the *SORL1*-202 reference transcript were observed in all three tested regions of the minipig brain for both the 5’-RT-PCR (exon 1-3, **Fig. 1c**) and 3’-RT-PCR (exon 46-47, data not shown) verifying the presence of exon 1 in pig *SORL1* transcripts.

### Generating *SORL1* gene edited Göttingen Minipigs

We used CRISPR-Cas9-mediated gene editing and cloning to create Göttingen Minipigs with compromised *SORL1* expression due to a heterozygous deletion of 609 bp including the entire *SORL1* exon 1 (**Fig. 1d**). Gene edited Göttingen Minipigs fibroblasts were used for cloning by somatic cell nuclear transfer (SCNT) to generate the *SORL1*-compromised animals (**Fig. 1e,f**).

SCNT resulted in *SORL1*^+/−^ animals as expected but also in delivery of a single piglet with homozygous knockout of *SORL1.* In contrast to the remaining piglets, this cloned (F0) *SORL1*^−/−^ piglet survived the post-natal period and served as a control in the study (from here on referred to as *ko)*. Re-cloning of fibroblasts obtained from one of the newborn cloned *SORL1*^+/−^ piglets resulted in delivery of live-born founder (F0) piglets without visible gross abnormalities and harboring the expected *SORL1*^+/−^ genotype (from here on referred to as *het*) as validated by Southern blot analysis and PCR-based genotyping (**Supplemental Fig. S1**). Two sexually mature *SORL1-het* female founders were used for conventional breeding with wild-type Göttingen Minipigs boars yielding two litters of naturally bred offspring (F1, 6 *het* and 4 *wt;* details on gender, genotypes, and ID numbers for all animals included in the study are provided in **Supplemental Table S1**).

To verify that the CRISPR-Cas9 gene editing strategy employed did not introduce unintended off-target effects, we analyzed genomic DNA isolated from both the re-cloned *het* piglets and a wild-type Göttingen Minipigs. Eight potential off-target sites residing in annotated genes were identified when allowing for up to three mismatches between the single-guide RNA (sgRNA) and the genomic sequence. These genomic sequences, comprising the potential sgRNA binding sites, were amplified by PCR and the resulting amplicons sequenced. We found no evidence of Cas9 induced off-target effects in any of the genomic sites analyzed, nor did we by PCR analyses observe random integration of the vectors employed (encoding the sgRNA and humanized Cas9, respectively) in the genomic DNA of the cloned *SORL1-het* and *SORL1-ko* animals (data not shown).

### Confirming *SORL1* haploinsufficiency in the *SORL1-het* minipig model

To verify that genomic deletion of *SORL1* exon 1 resulted in loss-of-function and reduced *SORL1* expression, reverse transcriptase PCR (RT-PCR) was performed on cDNA obtained from total RNA isolated from cerebellum (Cb) and frontal cortex (Cx) of brains from three sacrificed pigs (F1 *wt* (6478)/5 mo; F1 *het* (6473)/5 mo, and the cloned F0 *ko* (6304)/33 mo) using a primer pair spanning the exon 1-2 boundary. The analysis confirmed that our knock-out strategy had resulted in complete removal of exon 1 comprising *SORL1* transcripts in the cloned *ko* minipig. Further analysis using a primer pair spanning *SORL1* exon 46-47 revealed that no other *SORL1* transcripts were present in the *ko* sample, strongly indicating the lack of alternative ways to initiate transcription and that exon 1 is essential for *SORL1* transcription in these brain regions of pigs (**Fig. 2a**).

**Figure 2.**
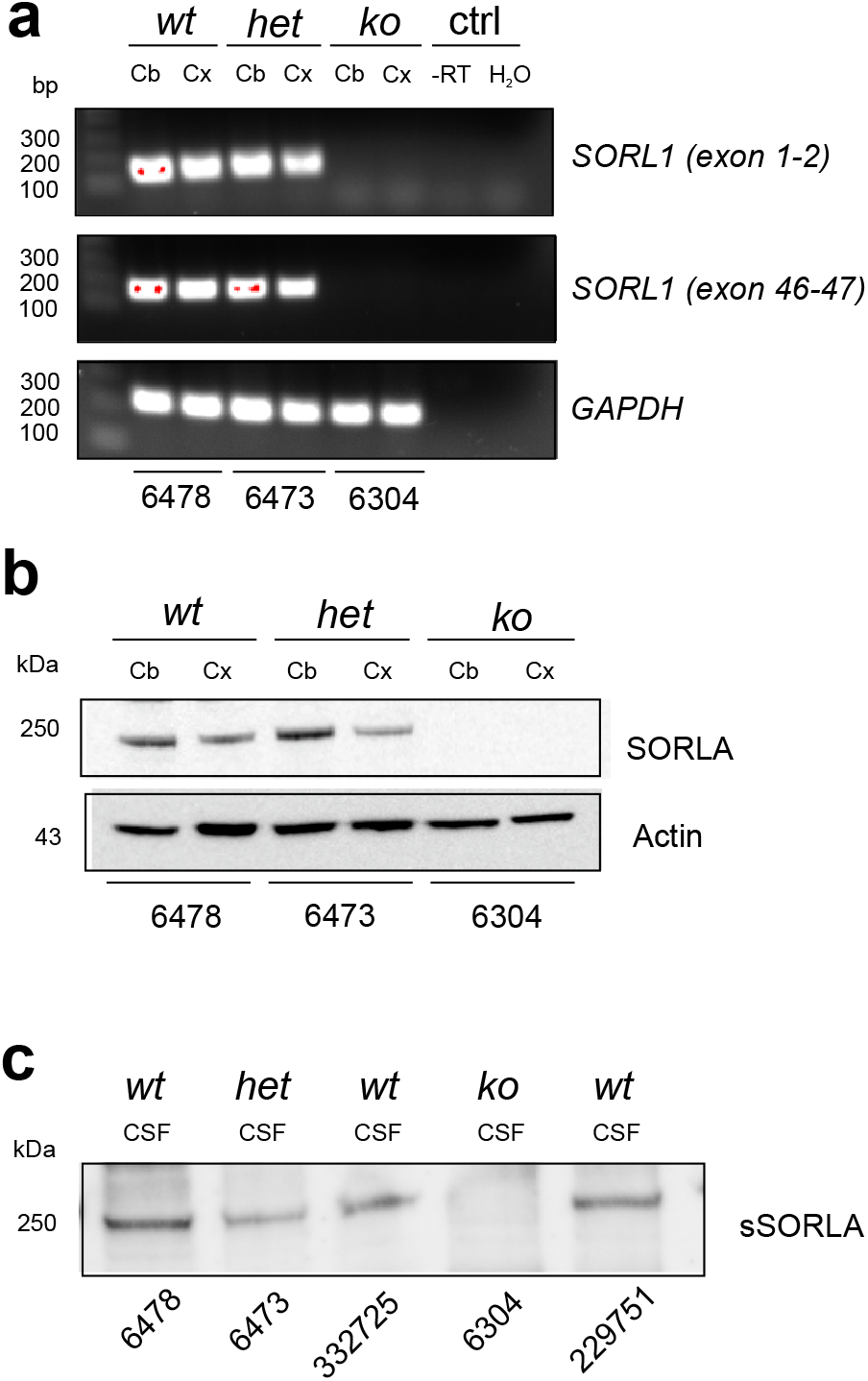
Decreased SORLA expression in brain of *SORL* 1-deficient Göttingen Minipigs. **a)** RT-PCR validation of *SORL1*-depleted Göttingen Minipigs. cDNA obtained from reverse transcription of total RNA isolated from either cerebellum (Cb) or frontal cortex (Cx) from *wt*, *het*, and *ko* Göttingen Minipigs was used as template in RT-PCR using primers specific for *SORL1* exons 1-2 (see primers F1+R2 in Fig. 1d), *SORL1* exons 46-47 of the *SORL1*-202 transcript, or for porcine GAPDH (reference gene). Negative controls (ctrl) correspond to reactions carried out in parallel using either RNA without reverse transcription (-RT) or with water (H_2_O) as templates. Western blot analysis of SORLA expression in cerebellum (Cb) or frontal cortex (Cx) homogenates (**b**) or cerebrospinal fluid (CSF) (**c**) from *wt*, *het* and *ko SORL1* Göttingen Minipigs. Identification numbers of individual pigs are provided below every lane.

Translation of the *SORL1-202* transcript is predicted to produce a SORLA protein of 2,213 amino acids that share 92.2% identity with the 2,214 residues of human SORLA (**Supplemental Fig. S2**). SORLA is a transmembrane protein but is also found in a soluble form, sSORLA, resulting from proteolysis and shedding from the neuronal cell surface (24, 25).

To corroborate our RT-PCR findings at the protein level, we next analyzed membrane-bound SORLA levels in homogenates of frontal cortex and cerebellum, isolated from brains of the three sacrificed minipigs (F1 *wt* (6478), F1 *het* (6473), and F0 *ko* (6304)) by Western blot (WB). As expected, SORLA could not be detected in samples from the *ko* minipig, whereas decreased expression of SORLA was observed in the cortex of the *het* minipig when compared to its *wt* littermate (**Fig. 2b**). Interestingly, the level of SORLA in the cerebellum was unaffected by the presence of only a single functional *SORL1* copy suggesting that SORLA levels can be maintained in the heterozygous state in cerebellum – a finding in line with it being a region free of most pathological defects and with intact SORLA expression in AD patients (26, 27). Likewise, WB analysis of cerebrospinal fluid (CSF) isolated from the same animals and two older control animals (*wt* (332725)/22 mo and *wt* (229751)/23 mo) confirmed absence of sSORLA in the *ko* animal, and showed that the *het* minipig has reduced SORLA expression reflected by lower sSORLA levels in the CSF (**Fig. 2c**).

As part of the validation of the model, neuropathological examination and histology was performed on brain sections from the frontal, temporal and occipital lobes, the brain stem and cerebellum obtained from a *het* (6473) and a *wt* (6478) F1 minipig at 5 months of age, respectively. Macroscopically, the ventricular system was not found to be dilated and no apparent brain pathology was observed in the cortex in these animals. Subsequent microscopical examination of hematoxylin/eosin-stained sections also revealed normal brain morphology including a normal ventricular ependymal layer, normal hippocampi and no neuron loss or neuron atrophy in either of the animals. (**Supplemental Fig. S3**).

From these analyses, we conclude that the *SORL1*-*het* minipigs have decreased expression of functional SORLA protein in cortex and CSF similar to AD patients harboring *SORL1* truncating mutations.

### Elevated CSF Aβ and tau in young *SORL1-het* minipigs

The sequence identity between human and porcine APP is 97.8% (**Supplemental Fig. S4**), and importantly, the 42 amino acids that correspond to the Aβ peptide is 100% conserved. Accordingly, we determined the level of the amyloid β-peptides (Aβ40 and Aβ42) in CSF isolated and combined from the cohort of our minipigs (*wt*, N=10; *het*, N=5; *ko*, N=1), using an available mesoscale discovery (MSD) assay developed for the human peptides. For analysis, samples from the *het* minipigs and the single *ko* animal were combined and compared to the group of *wt* minipigs. Despite that samples were collected from animals across different ages (*wt*, mean age = 22.4 ± 3.0 mo; *het+ko*; mean age = 23.2 ± 4.5 mo) we found a robust increase in the level of Aβ peptides (149 % for Aβ40 and 169 % for Aβ42) in CSF from *SORL1*-deficient (combined *het* and the single *ko*) minipigs (**Fig. 3a,b)**. Also, statistical analysis omitting the sample isolated from the *ko* minipig (data marked as red in figure 3) showed a significant (*P=0.0056* (Aβ40) and *P=0.0023* (Aβ42)) increase of CSF Aβ in the cohort of *SORL1-het* minipigs.

**Figure 3.**
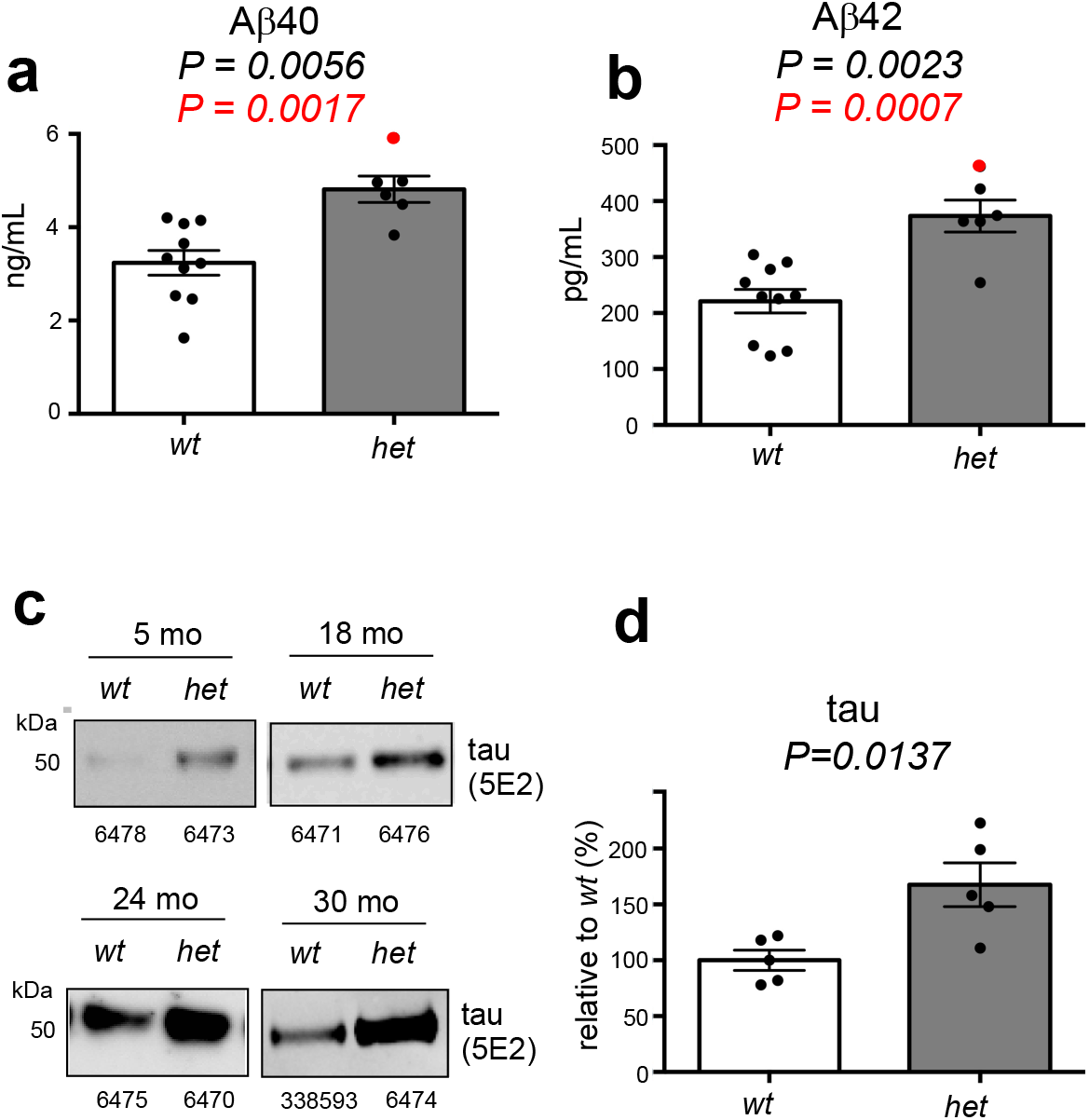
Increased APP and tau levels in CSF from *SORL1*-deficient Göttingen Minipigs. Quantification of APP processing products Aβ40 (**a**) and Aβ42 (**b**) in CSF from *SORL1*-deficient (N=6) and age-matched wildtype (N=10) Göttingen Minipigs. Average age of the two groups of pigs were similar. The group of *het* minipigs were depicted including data obtained from the *ko* pig 6304 shown in red, and statistical analysis shown for data excluding (black) or including (red) the data point for the *SORL1-ko* pig. Quantifications were performed using MSD assays for human APP fragments due to 100% conservation of the 42 amino acids comprising the Aβ sequence. **c)** Western blot analysis of tau in CSF (first time isolation) from *wt* and *het SORL1* minipigs at 5, 18, 24 or 30 months of age. Identification numbers of individual pigs are provided below every lane. Detection was performed with the 5E2 anti-tau antibody that binds to a region of tau that is 100% conserved between the human and pig protein (**Supplemental Figure S5**). **d)** Quantification of CSF tau Western blot analysis for the 18-month old (1 *wt*/*het* pair), 24-month old (2 *wt*/*het* pairs), and 30-month old (2 *wt*/*het* pairs) minipigs. The signal for *SORL1-wt* pigs were for each age set to 100% and the signal for the paired *SORL1-het* expressed relative to this.

We next analyzed tau levels in CSF from pairs of age-matched *het* and *wt* minipigs with CSF isolated at 5, 18, 24 or 30 months of age (**Fig. 3c**). As the sequence homology between porcine and human tau (89.8% sequence identity; **Supplemental Fig. S5**) varies in the region utilized in standard assays for tau detection, WB analysis using an antibody (5E2) for tau mid-fragment (~50 kDa) was applied to test CSF from 5-30 month old minipigs (**Fig. 3c**) and quantify total-tau levels in CSF from the 18-30 month old minipigs (*wt*, N=5; *het*, N=5) (**Fig. 3d**).

In all four comparisons, elevated ~50 kDa tau levels were observed in the *SORL1-het* samples (**Fig. 3c**), and the tau levels were found to be significantly increased when analyzing the samples as groups irrespective of age (*P*=0.0137, **Fig. 3d**).

These findings of elevated levels of both Aβ and tau in the CSF of the *SORL1-het* minipigs follow observations of increased amyloid and tau pathology in cellular models of endosomal dysfunction (28, 29), and are in accordance with previous findings of increased levels of CSF Aβ in the very early phase of disease progression in sporadic AD patients (3), patients with autosomal dominant AD (4), and in transgenic AD mouse models prior to plaque formation (5).

### 21-months old *SORL1-het* minipigs do not exhibit amyloid plaque formation

We next employed positron emission tomography (PET) imaging with the tracer [^11^C] *N*-methyl [^11^C] 2-(4′methylaminophenyl)-6-hydroxy-benzothiazole (PIB), which allows for visual and quantitative measurement of Aβ deposition, to analyze if amyloid plaques had started to form in the brains of the *SORL1-het* minipigs. We subjected four 21-month old female minipigs (*het* (N=2) and *wt* (N=2)) to [^11^C]-PIB-PET imaging, but we did not observe any obvious increase in tracer retention in the time activity curves of the *het* compared to the *wt* animals suggesting that fibrillar Aβ is not yet accumulating in the brains of these young *SORL1-het* animals (**Fig. 4a**), in agreement with the measured increase rather than decrease in CSF Aβ.

**Figure 4.**
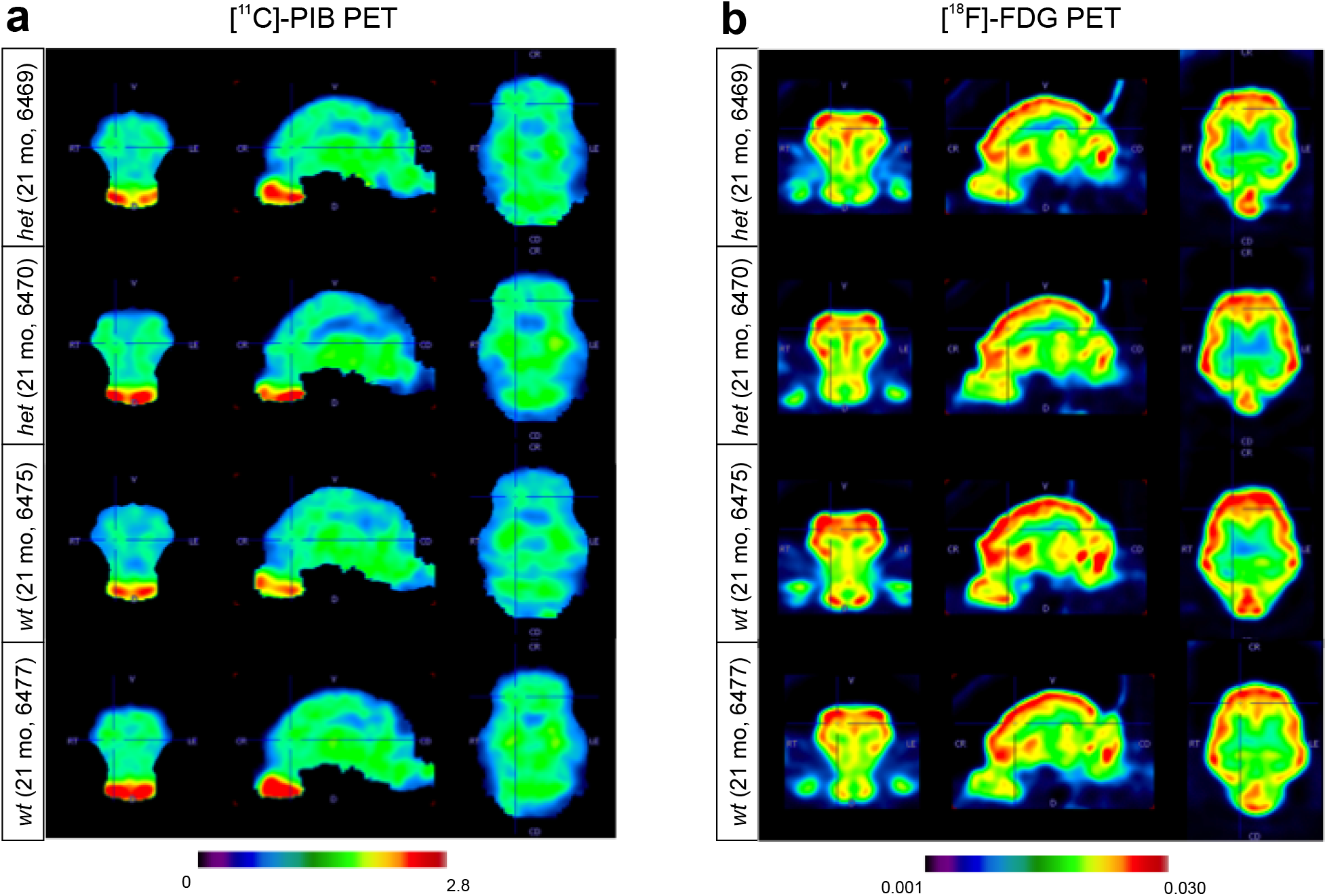
[^11^C]-PIB and [^18^F]-FDG PET analysis of young *SORL1-het* and *SORL1-wt* Göttingen Minipigs. [^11^C]-PIB- and [^18^F]-FDG PET analyses of four female 21-month old minipigs (*het*: 6469 and 6470; *wt*: 6475 and 6477). **a)** [^11^C]-PIB-PET summed images from the 30-90 minute portion of the dynamic scan, divided by averaged whole brain activity. **b)** [^18^F]-FDG PET standard uptake volume (SUV) images, corrected for full brain activity.

We also investigated brain hypometabolism as a marker of neurodegeneration in the four 21-month old female minipigs (*het* (N=2) and *wt* (N=2)) by PET imaging with 2-[^18^F]-2-deoxy-D-glucose (FDG) which measures the cerebral metabolic rates of glucose as an indicator of neuronal synaptic activity (30). We did not observe any decrease of [^18^F]-FDG-tracer uptake in the brains of the *SORL1-het* minipigs (**Fig. 4b**). This suggests that any pathological changes occurring in the brain of these young *SORL1-het* minipigs are not sufficient to cause hypometabolism reflective of neuronal degeneration at this stage.

### *SORL1-het* minipigs display unaltered brain morphology and neuroaxonal damage up to 27 months of age

Having established that the observed CSF changes in the *SORL1-het* minipigs were not accompanied by amyloid deposition and brain hypometabolism, we examined if the animals exhibited changes in brain morphology by subjecting the now 22-month old four female minipigs (*wt*, N=2; *het*, N=2) as well as four 27-month old males (*wt*, N=2; *het,* N=2) to anatomical magnetic resonance imaging (MRI). We observed variation between animals, mainly between males and females, but were not able to detect any sign of genotype-dependent brain atrophy in these young *SORL1-het* minipigs, including any difference in overall brain volume (**Fig. 5a,b**). Likewise, individual brain structures considered to be the first affected by atrophy in AD, did not show statistically significant differences between *SORL1-wt* and *SORL1-het* minipigs as illustrated by the measurement of amygdala width, entorhinal cortical thickness, and hippocampal and ventricular volumes (**Fig. 5c–f**).

**Figure 5.**
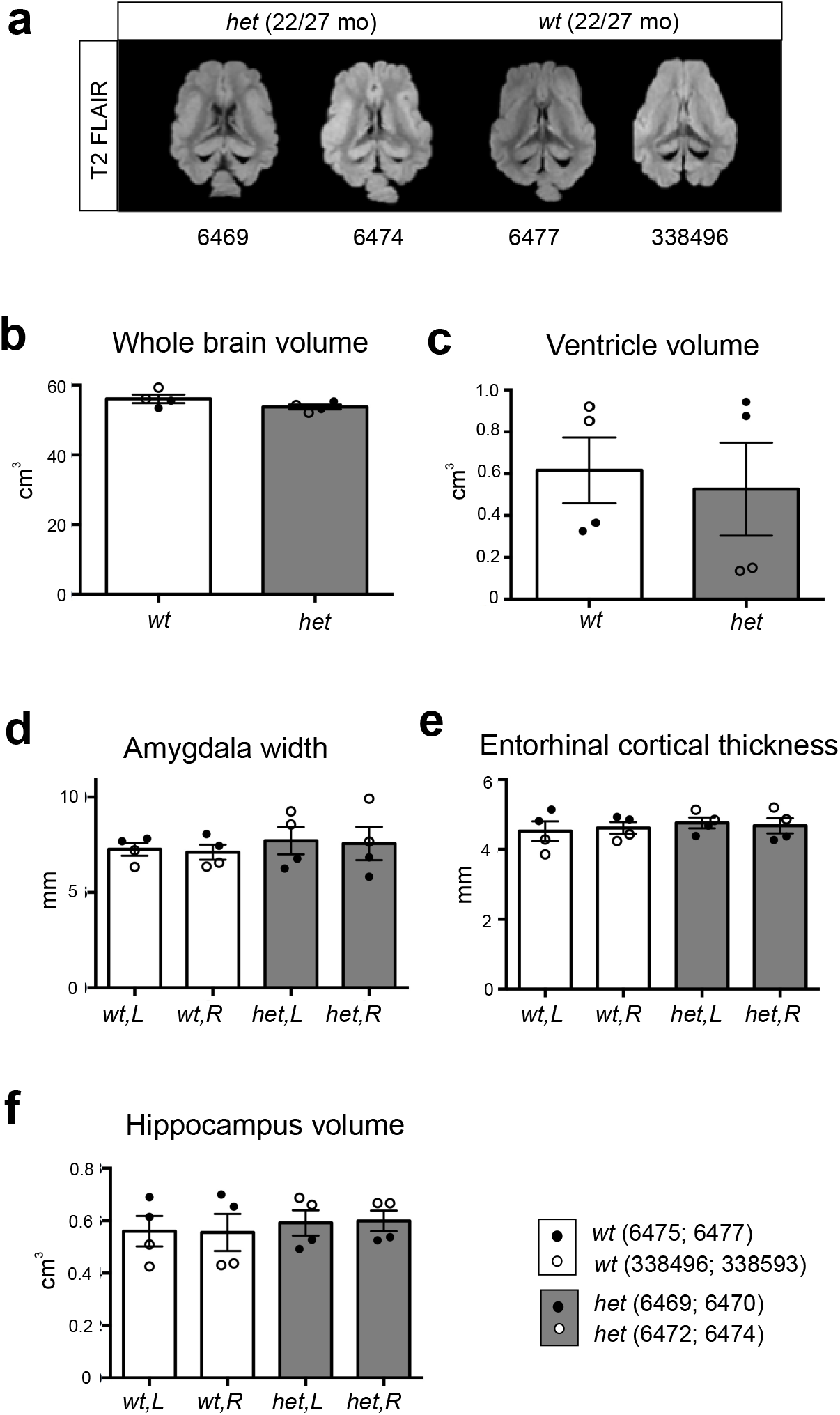
Anatomical MRI analysis of young *SORL1-het* and *SORL1-wt* Göttingen Minipigs. **(a-f)** Examples of 3D T2 FLAIR magnetic resonance images used for volume measurements (extracerebral tissue stripped for visualization) in female (6469) and male (6474) *SORL1-het* minipigs and female (6477) and male (338496) *SORL1-wt* minipigs (**a**). Quantification of whole brain volume (**b**), ventricle volume (**c**), amygdala width (**d**), entorhinal cortical thickness (**e**) and hippocampus volume (**f**) from anatomical MRI-scanning. The four female minipigs (*wt*: 6475 and 6477; *het*: 6469 and 6470) and the four male minipigs (*wt*: 338496 and 338593; *het*: 6472 and 6474) were 22 and 27 months of age when scanned, respectively.

Having demonstrated normal brain anatomy with anatomical MRI, we next determined if there were more subtle structural changes in brain microstructure using diffusion tensor imaging (DTI), a more sensitive method for detecting microscopic damage than conventional structural MRI (31) enabling detection of potential diffusivity changes due to cell loss, demyelination and axonal injury. We subjected the eight minipigs to DTI in connection with the structural MRI scan but found no changes in mean diffusivity (MD) or fractional anisotropy (FA) in the amygdala, hippocampus, corpus callosum and deep white matter between minipigs of the two genotypes (**Fig. 6a–e**).

**Figure 6.**
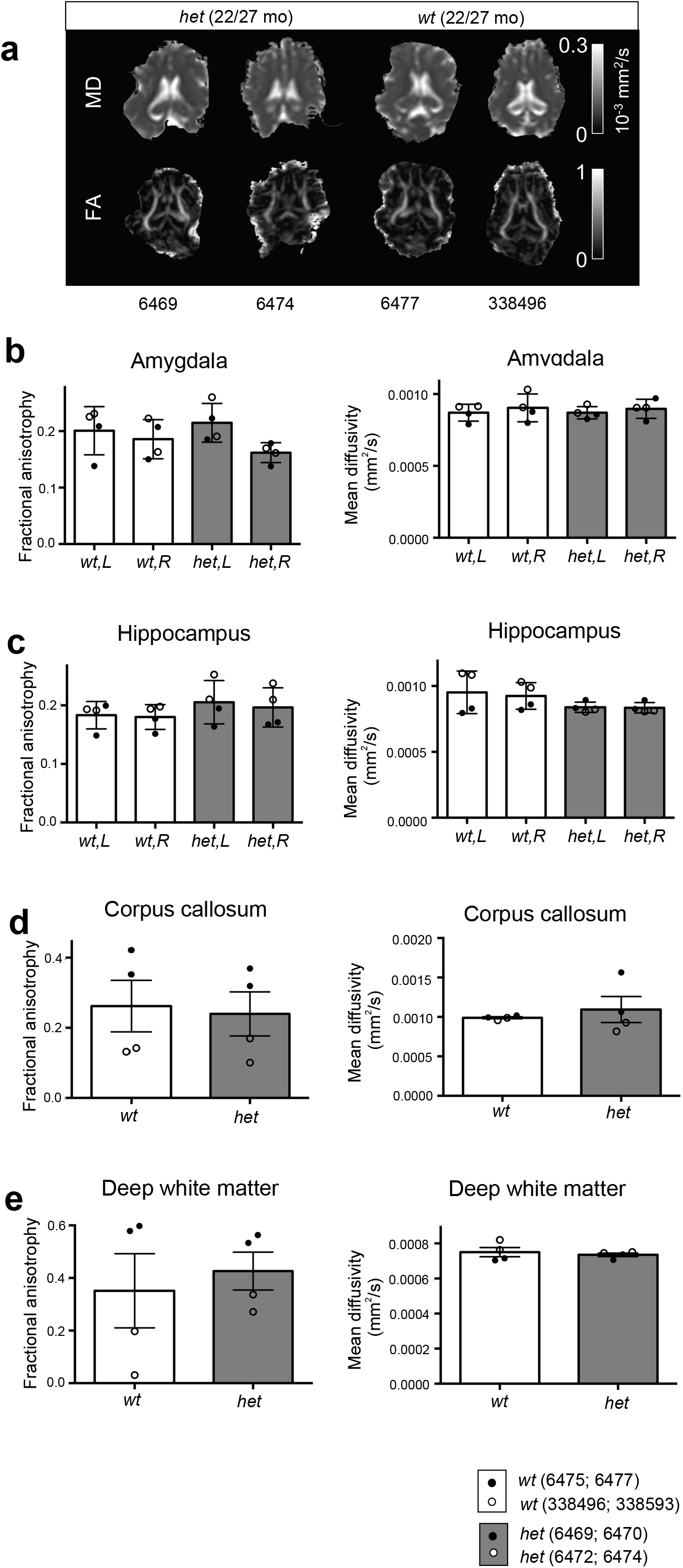
Normal brain microstructure in *SORL1-het* and *SORL1-wt* Göttingen Minipigs. Examples of mean diffusivity (MD) and fractional anisotropy (FA) maps, representing tissue microstructure, from diffusion tensor MRI in female (6469) and male (6474) *SORL1-het* minipigs and female (6477) and male (338496) *SORL1-wt* minipigs (**a**). Fractional anisotropy (FA) and mean diffusivity (MD) of the amygdala (**b**), hippocampus (**c**), corpus callosum (**d**) and deep white matter (**e**) were determined with diffusion tensor imaging. The four female minipigs (*wt*: 6475 and 6477; *het*: 6469 and 6470) and the four male minipigs (*wt*: 338496 and 338593; *het*: 6472 and 6474) were 22 and 27 months of age, respectively, when scanned.

Finally, we quantified the level of Neurofilament light chain (NF-L), a biomarker indicative of axonal damage. NF-L is used for monitoring progression in many neurological disorders and has been suggested for tracking neuronal fitness during AD (32). In the CSF that was collected from minipigs at necropsy (*wt*, N=10; *het*, N=5; *ko*, N=1), and also used for Aβ40/42 analyses, we did not detect any significant difference between *SORL1-het* and *SORL1-wt* minipigs for this biomarker (**Supplemental Fig. S6**).

Collectively, our results suggest that the 5-30 month old *SORL1-het* minipigs are in an early preclinical phase of AD with raised CSF Aβ and tau levels prior to progression to the more advanced AD pathology including amyloid plaque formation and neuronal loss. Our *SORL1-het* minipig model thus phenocopies the preclinical *in vivo* profile of AD observed with other established AD causal genes (4).

## DISCUSSION

Since genetic epidemiology has suggested that the most pathogenic *SORL1* variants harbor truncating mutations that cause a partial loss of protein function, modelling this pathogenic state can be achieved by inducing *SORL1* deficiency. Here we have by CRISPR-Cas9-mediated gene editing developed a Göttingen Minipigs model with *SORL1* haploinsufficiency. Previous work employing mouse models and neuronal cultures have suggested that *SORL1* deficiency phenocopies the cell biology of the established AD-causal genes encoding APP and the presenilins. Just as seen with these causal genes, *SORL1* deficiency accelerates amyloid production in the early endosome of neurons (17, 33), also causing endosomal traffic jams which manifest as swollen neuronal endosomes (19), a hallmark feature of AD neurons (34).

In both the case of APP and the presenilins, their mutations accelerate the endosomal accumulation of the intermediary APP fragment β-CTF, the direct precursor of Aβ, and it is this membrane-spanning fragment, not Aβ, that is suspected to cause endosomal traffic jams (6, 35). In the presented *SORL1* depleted minipigs, endosomal traffic jams relate to SORLA acting as a receptor for recycling cargo out of the endosome. Both APP and β-CTF are cargo for SORLA-dependent endosomal recycling (17, 36, 37). The decreased SORLA endosomal activity therefore leads to an increased Aβ production, that manifest as elevated CSF Aβ levels.

While these cell biological investigations are important in trying to elucidate mechanisms of these genes, they by necessity were not, and cannot, be confirmed in actual patients. We have previously shown that complete deletion of *SORL1* in mice leads to increased amyloidogenesis, nevertheless no analysis of CSF biomarkers were performed during the course of disease progression as sampling of CSF to monitor e.g. early changes in biomarkers is a challenging procedure in mice thus hampering detection of early biochemical pathology in this animal model (17, 33, 38). The current study employing gene-edited Göttingen Minipigs with *SORL1* haploinsufficency, mimicking the genetic status of AD patients with *SORL1* haploinsufficiency, is thus the first to show an *in vivo* phenocopy of *SORL1* deficiency to AD’s causal genes, observed in actual patients in the earliest preclinical phase of the disease (i.e. elevated Aβ and tau in CSF prior to formation of amyloid plaques and neurodegeneration). While cell biological observations cannot be confirmed in patients, their mechanistic implications can help explain the *in vivo* profile observed. Only through cell biology do we know, for example, that amyloid is produced and first accumulates inside neuronal endosomes (7), when APP is cleaved by its amyloidogenic enzymes at the endosomal membranes, and that intraneuronal amyloid is then secreted unconventionally by endosomal secretion (39). *AP*P and *PSEN* mutations are known to accelerate amyloid production by directly affecting the amyloidogenic biochemical pathway. *SORL1* deficiency, in contrast, indirectly leads to accelerated amyloid production by slowing APP’s recycling out of endosomes, thereby increasing the probability that it will be cleaved at endosomal membranes harboring high β-secretase activity (40), and results in an increase of Aβ peptides in CSF of both 40 and 42 amino acids in length.

More recent cell biological studies have clarified why tau accumulates in the CSF in the setting of mutations in APP and the presenilins. It is now known that tau is actively secreted from neurons, and just like amyloid, occurs in CSF via endosomal secretion (41, 42). The abnormal elevation of CSF tau observed in AD is no longer thought to be driven by tangle formation or neurodegeneration per se, but is rather an early event that reflects accelerated active secretion, one that seems to coincide with amyloid secretion (43). The precise details of tau’s accelerated secretion remain unknown, but likely relate to how endosomal traffic jams leads to increased translocation of tau into the endolysosomal system. Since the endosomal traffic jams induced by defects in APP, the presenilins, or SORLA are shown to be amyloid β-peptide independent, amyloid and tau secretion are likely coincidental to each other.

In summary, by showing that our gene-edited *SORL1* haploinsufficient Göttingen Minipigs phenocopy the biomarker profile of the earliest preclinical stage of disease observed in patients carrying known causal mutations, we provide functional evidence that *SORL1* loss-of-function mutations are causal in AD. After exhaustive and large-scale genetic investigations into AD, it seems likely that there are only four genes across our genome that can themselves cause AD: *APP, PSEN1, PSEN2*, and *SORL1*. Moreover, interpreting the unified *in vivo* profile induced by these four causal genes in the context of their cell biological consequences, suggests that dysfunctional endosomal trafficking is a unified pathogenic pathway in the disease that could be pursued for drug development (16, 44).

Besides providing evidence that *SORL1* is an AD-causing gene, the *SORL1-het* Göttingen Minipigs allow for future longitudinal studies for novel biochemical and neuroimaging biomarker discovery and may thereby provide important clues to the sequence of events that occur in the pharmacologically valuable treatment window between the very early preclinical stage of the disease, with raised Aβ and tau CSF levels, and the time point of amyloid brain deposition, irreversible neurodegeneration, and subsequent cognitive impairment.

Due to their similarities to humans in terms of genetics, anatomy, physiology and biochemistry, minipigs are in addition valuable animal models for drug testing and are being increasingly employed as non-rodent models for toxicity testing and safety pharmacology (45, 46) thus adding to the utility of our Göttingen Minipigs *SORL1-het* model in pharmacological research targeting AD.

## METHODS

### Animals

Permission for generation and breeding as well as for neuroimaging and blood/CSF sampling of the *SORL1* Göttingen Minipigs were granted by the Danish Animal inspectorate (2015-15-0202-00028 and 2019-15-0201-00264). Göttingen Minipigs were housed under standard conditions in specific pathogen-free housing facilities at Aarhus University and maintained on a restricted minipig diet (SDS Diet, UK) throughout the study. The minipigs were all acclimatized for at least one week prior to experiments and housed, in groups after weaning until 4 months of age and thereafter as single animals, under environmental conditions of 20-22-C, 50–55% relative humidity, 12:12 hours of light and darkness, and with change of air on at least eight times per hour. They had free access to tap water, and their well-being was monitored twice per day. The welfare of the animals was improved by access to “easy-strø” bedding (Easy-AgriCare A/S, Denmark) and chewing toys (e.g. sisal ropes).

### Analysis of porcine SORL1 transcripts

Total RNA was isolated from cortex, hippocampus and cerebellum from wildtype Göttingen Minipigs using an RNeasy Micro Kit (#74004, Qiagen). cDNA was produced using 1 μg of total RNA and a RevertAid First Strand cDNA Synthesis Kit (#K1622, Thermo Fischer Scientific) prior to performing RT-PCR analyses to validate the presence of the reference Sscrofa 11.1 *SORL1*-202 transcript. RT-PCR analyses were performed using GoTaq DNA polymerase (#M7841, Promega) in a reaction volume of 25 μl comprising 1 μl of cDNA and primer pairs specific for the 5’-end (exon 1-3, see primers F1+R1 in Fig. 1d) and 3’end (exon 46-47), respectively, of the *SORL1*-202 transcript or with primers specific for the reference gene *GAPDH*. The following PCR conditions were employed: 95°C for 2 mins for 1 cycle followed by 35 cycles of 95°C for 30 s; 58°C/60°C/65°C (depending on the primer set) for 30 s, 72°C for 30 s followed by 1 cycle of 72°C for 7 mins. The resulting *SORL1-202* amplicons were spin-column purified using a Nucleospin Gel and PCR Clean-up Kit (#740609, Macherey-Nagel) prior to validation by DNA sequencing. Sequences of primers used for the RT-PCR analyses are provided in **Supplemental Table S2**.

### Cell culture

Primary porcine fibroblasts were established by explant culture of ear biopsies from newborn Göttingen Minipigs obtained from Ellegaard Göttingen Minipigs A/S. Until fibroblast outgrowth, ear biopsies were cultured in AmnioMAX-C100 complete medium (#12558-011, Thermo Fischer Scientific). Upon isolation, the porcine fibroblasts were cultured at 37°C in DMEM (#BE12-604F, Lonza Biowhittaker) supplemented with 15% heat-inactivated fetal calf serum, 60 μg/mL penicillin, 100 μg/mL streptomycin, and 292 μg/mL glutamine. The cell culture medium was additionally supplemented with recombinant basic fibroblast growth factor (bFGF, 5 ng/mL, #PHG0266, Thermo Fischer Scientific) for gene targeting experiments.

### Construction of gene targeting vector and single-guide RNAs

A gene targeting vector was generated essentially as previously described (47). The left and right porcine *SORL1* (p*SORL1*) homology arms (LHA and RHA, 880 bp and 998 bp, respectively) flanking *SORL1* exon 1 were amplified by PCR using wild type Göttingen Minipigs genomic DNA as template and porcine *SORL1*-specific primers. These two homology arms were subsequently linked to a neomycin (*neo*)/zeomycin (*zeo*) resistance gene cassette allowing for G418 selection by a three-way fusion PCR. The *neo*/*zeo* resistance genes were comprised in a 4 kb *PvuI* fragment isolated from a pNeDaKO-Neo plasmid (a generous gift from Bert Vogelstein & Kenneth W. Kinzler, The Johns Hopkins University Medical Institutions, Baltimore, MD 21231, USA). The 3-way fusion PCR was performed as described (47) using a Platinum Pfx polymerase (#11708-013, Thermo Fischer Scientific) and the following PCR protocol: 1 cycle of 94°C for 1 min; 25 cycles of 94°C for 30 s, 59°C for 30 s, and 68°C for 4 min; 1 cycle of 68°C for 7 min. Fusion products were digested with *NotI* and ligated to a *NotI* cleaved pAAV-MCS plasmid backbone (#240071-5, Stratagene). The final rAAV/*SORL1* KO-Neo plasmid construct was verified by *NotI* digestions and sequencing. Successful gene targeting using the rAAV/*SORL1* KO-Neo plasmid vector results in deletion of 609 bp in the endogenous porcine *SORL1* gene including the entire exon 1.

The CRISPR/Cas9 single-guide RNAs (sgRNAs) targeting the porcine *SORL1* gene were designed employing the online designing tool ZiFiT (http://zifit.partners.org/ZiFiT/). The human codon-optimized Cas9 (kindly made available by George Church, Addgene plasmid # 41815) and the sgRNA (encoded by a pFUS-U6 vector) were expressed from two individual plasmids. Two different sgRNAs, sgRNA 1 and sgRNA2, were designed to target the porcine *SORL1* exon 1 region. For each sgRNA construct, two p*SORL1*-specific complementary oligonucleotides were denatured and slowly annealed prior to ligation of the annealed oligonucleotides to a sgRNA scaffold plasmid (pFUS-U6-sgRNA) based on *Bsa*I assembly as previously described (48). XL-2 Blue ultracompetent bacterial cells (#200150, Agilent Technologies) were subsequently transformed with the ligation mixture and the resulting bacterial cell clones were screened by PCR. Positive sgRNA clones were validated by DNA sequencing of purified plasmid DNA. PCR primers for generating the rAAV/*SORL1* KO targeting vector as well as oligonucleotides and target sites used for generation of the sgRNA vectors are listed in **Supplemental Table S2.**

### Generation of pSORL1-specific C-check vector for sgRNA testing

The two p*SORL1*-specific sgRNAs, sgRNA1 and sgRNA2, were functionally validated employing a previously developed single strand annealing (SSA)-directed, dual fluorescent surrogate reporter system entitled C-check (48). This vector comprises two expression cassettes: an *AsRED* expression cassette for measuring transfection efficiency and normalization, and a truncated *EGFP* expression cassette for detection of double strand break (DSB)-induced SSA events. The *EGFP* cassette is interrupted by the two target sites for the p*SORL1* sgRNAs. The C-check reporter construct will express *AsRED* and *EGFP* upon sgRNA-induced DSB and SSA repair in this target region, whereas only *AsRED* will be expressed if no DSB is induced, or repair occurs by non-homologous end joining. To construct the p*SORL1*-specific C-check vector, two complementary oligonucleotides comprising the sgRNA target sites were annealed and cloned into the *Bsa*I-digested C-check vector as previously described (48). XL-2 Blue Ultracompetent cells were transformed with the ligated plasmid, and resulting bacterial cell clones were screened by PCR. Positive C-check clones were validated by DNA sequencing upon purification of plasmid DNA. The oligonucleotides employed to construct the p*SORL1*-specific C-check vector comprising the overlapping sgRNA1 and sgRNA2 target sites are listed in **Supplemental Table S2**.

### Flow cytometric analysis of pSORL1 gRNA efficiency

The efficiencies of the two generated p*SORL1*-specific sgRNAs were evaluated by transfection of the sgRNA-encoding plasmid into HEK293T cells and subsequent flow cytometry of the transfected cells as previously described (48). Briefly, cells were seeded into 6-well plates (3 ×10^5^ cells/well) and co-transfected the next day with one of the two sgRNA plasmids (75, 150, and 300 ng) together with the hCas9 plasmid (75, 150, and 300 ng), the p*SORL1-*specific C-check plasmid (100, 200, and 300 ng), and stuffer plasmid DNA (to adjust the amount of total DNA to 1 μg) using X-tremeGENE 9 DNA transfection reagent (#6365779001, Sigma-Aldrich). For controls, cells were transfected with only the hCas9 or C-check plasmid, or with the sgRNA construct together with the C-check plasmid. Cells were harvested 48 hours post transfection by trypsinization and analyzed by flow cytometry using a BD LSRFortessa Flow Cytometer (BD Bioscience) at the FACS Core Facility, Dept. of Biomedicine, Aarhus University, to quantify the efficiency (*EGFP* expression) of the two sgRNAs in transfected (*AsRED*+) cells. Based on these results, sgRNA1 was chosen for CRISPR-based gene editing of primary Göttingen Minipigs fibroblasts.

### Gene editing

The day before transfection, primary fibroblasts isolated from newborn female Göttingen Minipigs were seeded (1.5 × 10^6^) into a gelatin-coated 10 cm cell culture dish. The next day, the culture medium was changed and supplemented with bFGF (5 ng/μL) and gene editing was performed by co-transfecting the cells with the gene targeting rAAV/*SORL1* KO-Neo vector (5600 ng), the hCas9 plasmid (1200 ng) and the sgRNA1 vector (1200 ng) using Lipofectamin LTX Reagent (#15338500, Thermo Fischer Scientific). The transfected cells were trypsinized 48 hours post transfection, and ½ of the cell suspension was subjected to limiting dilution by reseeding the cells into 5 gelatin-coated 96-well plates resulting in approx. 300 cells per well. Selection with G418 (0.8 mg/ml, #ant-gn, Invivogen) was initiated the following day and continued for two weeks. Following selection, the G418-resistant cell clones were trypsinized, 1/3 of the resulting cell suspension was transferred to 96-well PCR plates for PCR screening, and 1/3 was cultured in gelatin-coated 96-well cell culture plates for Southern blot analysis. The remaining 1/3 of the cell suspension was cultured in gelatin-coated 96-well plates, frozen at early passages and subsequently used as nuclear donor cells for SCNT.

### PCR screening of gene edited donor cells

PCR screening for successful *SORL1* gene targeting was performed on lysates of individual G418-resistant cell clones with primer pairs F3+R3 and F4+R4 amplifying the 5’ KO - and 3’ KO regions, respectively (see position of primers and primer sequences in **Fig. 1d** and **Supplemental Table S2**, respectively). First, G418-resistant cells in the 96-well PCR plates were centrifuged and re-suspended in 25 μL lysis buffer (50 mM KCl, 1.5 mM MgCl_2_, 10 mM Tris-Cl, pH 8.5, 0.5% Nonidet P40, 0.5% Tween, 400 μg/ml Proteinase K) (McCreath et al, 2000) prior to lysis (65°C for 30 min, 95°C for 10 min). The lysate (3 μL) was subsequently used as template in a PCR screening using a Platinum Pfx DNA polymerase (#11708-013, Thermo Fischer Scientific) using the following PCR conditions: (1) *SORL1* 5’ KO screening: 1 cycle of 94°C for 2 min, 35 cycles of 94°C for 20 s, 63°C for 30 s, and 68°C for 1.5 min followed by 1 cycle of 68°C for 7 min; (2) *SORL1* 3’ KO screening: 1 cycle of 94°C for 2 min, 35 cycles of 94°C for 20 s, 56°C for 30 s, and 68°C for 1 min followed by 1 cycle of 68°C for 7 min. Primers used for the *SORL1* gene targeting screening are listed in **Supplemental Table S2.**

### Southern blot analysis

The gene edited *SORL1*^+/−^ donor cells and cloned piglets were further validated by Southern blotting using a porcine *SORL1*-specific probe (887 bp) located upstream of the targeted region. Also, a *neo^r^*-specific probe (1162 bp), detecting the *neo^r^* encoding cassette in the targeting vector, was used to examine if the gene edited cell clones and resulting cloned piglets also harbored additional unwanted random integrations of the vector (see **Supplemental Fig. S1a**). Both Southern blot probes were generated by standard PCR and subjected to random labelling using a Prime-It II Random Primer Labelling Kit (#300385, Agilent) according to the manufacturer’s instructions. Genomic DNA (15 μg) isolated from cultured gene edited fibroblasts, or from ear biopsies taken from new-born piglets, was digested with *BlpI* restriction enzyme overnight. The digested samples were subjected to gel electrophoresis on a 0.7% agarose gel followed by vacuum blotting onto a nitrocellulose membrane. Pre-hybridization, and hybridization with the individual probes, were carried out at 42°C and all washing procedures were performed at 53°C. Primers for generating the *SORL1* and *neo^r^* probes, respectively, are listed in **Supplemental Table S2**.

### Cloning and embryo transfer

Two of the validated *SORL1* 3’ KO/*SORL1* 5’ KO gene edited cell clones were used as nuclear donor cells for somatic cell nuclear transfer (SCNT) by handmade cloning as described by (49). Cumulus-oocyte complexes harvested from slaughterhouse-derived sow ovaries were in-vitro matured and treated to remove cumulus cells and partially zonae pellucidae. The oocytes were bisected manually, and the cytoplasts without chromatin were collected. Each cytoplast was first attached to one nuclear donor cell before being fused (BTX microslide 0.5mm fusion chamber, model 450; BTX San Diego, US). After 1 h of incubation, each cytoplast-donor cell pair was fused with an additional cytoplast creating the reconstructed embryo. All reconstructed embryos were then incubated in culture medium for 5-6 days after which the blastocysts and morulae were selected based on morphology. Two pools of cloned gene edited embryos were prepared, and 82 and 90 blastocysts/morulae, respectively, were transferred surgically into two recipient landrace surrogate sows (50). Pregnancy was diagnosed in both sows by ultrasonography after approx. 25 days. Farrowing was hormonally initiated at day 114 by intra-muscularly injected prostaglandin (Estrumate, 2 ml/sow), and the sows farrowed totally 8 live piglets (7 and 1, respectively). All piglets, apart from one, died post-natally or over the next few weeks. The surviving piglet was genotyped as *SORL1*^−/−^ (*SORL1*-*ko*) demonstrating that the donor cell clone used for SCNT was not derived from a single gene targeted cell. Further examination of genomic DNA isolated from this *SORL1-ko* piglet showed, an on-target, but only partial, gene targeting obstructing one allele, whereas a large deletion was found on the other allele possibly induced by non-homologous end-joining of the free DNA ends resulting from CRISPR-mediated double-strand cleavage.

Fibroblasts were isolated from ear biopsies taken on the day of birth from all cloned piglets. *SORL1*^+/−^ fibroblasts isolated from one of these piglets were used for re-cloning upon sequence validation following the same protocol as described above. Sixty-eight and 69 re-cloned embryos were transferred to two surrogate sows, respectively. Both were diagnosed pregnant, but one aborted later. The remaining pregnant sow gave birth to 9 piglets, of which 6 were alive. These cloned (F0) piglets were genotyped as *SORL1*^+/−^ by PCR and gene targeting was validated by Southern blotting (**Supplemental Fig. S1a,b**). Two of these 6 cloned founder piglets survived the post-natal period and were, upon sexual maturity, mated with wild-type Göttingen Minipigs boars (referred to as “breeding boars” in **Supplemental Table S1**) resulting in two naturally bred F1 litters of *SORL1*^+/−^ and *SORL1*^+/+^ piglets (*het*, N=6 and *wt*, N=4 in total). The surviving cloned (F0) *SORL1-ko* minipig was included as a control in the study. The 2 breeding boars and the 4 F1 *wt* animals were used as controls in addition to 6 naturally bred wild-type Göttingen Minipigs obtained from Ellegaard Göttingen Minipigs A/S.

### Genotyping

Genotyping of both cloned and naturally bred Göttingen Minipigs *SORL1*^+/−^ and *SORL1*^+/+^ piglets were performed by standard PCR using a Platinum Superfi DNA polymerase (#12351010, Thermo Fischer Scientific) and primer sets for detecting the 5’- and 3’-*SORL1* KO region, respectively, in addition to a primer set detecting the wild type *SORL1* gene. The following PCR conditions were employed: 1 cycle of 98°C for 30 secs, 35 cycles of 98°C for 10 s, 60-66°C (depending on the primer set) for 10 s, and 72°C for 30 secs followed by 1 cycle of 72°C for 7 min. Primers utilized for genotyping are listed in **Supplemental Table S2**.

### CRISPR off-target analysis

The online CRISPR RGEN Cas-OFFinder algorithm was used for identifying potential off-target sites for the employed p*SORL1* sgRNA1. In addition to the targeted *pSORL1* gene, eight potential off-target sites residing in annotated genes on chromosomes 2 (*JUNB* and *ARHGAP26)*, 5 *(XRCC6)*, 6 *(GSE1)*, 8 (*PCDH7*), 9 (*HEPACAM*), 14 (*TXNRD2*) and 15 (*TWIST2*) were identified when allowing for up to 3 mismatches between the sgRNA and genomic sequence. These potential off-target regions were amplified by standard PCR using genomic DNA isolated from wild type or cloned *SORL1*^+/−^ Göttingen Minipigs, Platinum Pfx DNA polymerase (#11708-013, Thermo Fischer Scientific), and primer pairs for the specific genomic region. The resulting amplicons, comprising the sequence region surrounding the sgRNA binding site, were purified using a Nucleospin Gel and PCR Clean-up kit (#740609, Macherey-Nagel) and subjected to DNA sequencing to verify if off-target activity had occurred.

Genomic DNA isolated from cloned *SORL1*^+/−^ Göttingen Minipigs was in addition analyzed for potential unwanted random integration of the plasmid constructs used for co-transfection (sgRNA, hCas9) by standard PCR using primer sets specific for the individual plasmids used for transfection. Primers used for off-target and random integration analyses are shown in **Supplemental Table S2.**

### Sampling of cerebrospinal fluid, plasma and tissues

Minipigs were anaesthetized with Zoletil-mix (1 ml/10 kg) and blood samples were taken from the jugular vein for plasma preparations. The animals were then placed in sternal recumbency with the neck flexed, and the relevant part of the neck region was surgically prepared. The anatomical landmarks were the occipital protuberance and the two lateral sides of the atlas wings. A spinal needle (BD 20 Gauge 3.50 in.) was passed perpendically down to the atlanto-occipital intervertebral space and stopped when a weak reflex from the minipig was felt. The minipig was then carefully turned to right lateral recumbency and the cerebrospinal fluid (CSF) was collected by gravity and capillary action. Apart for 4 female F1 minipigs (6469, 6470, 6475, and 6477), which were subjected to CSF sampling twice, all minipigs were euthanized immediately after CSF sampling with an intravenous injection of 30% pentobarbital (0.25 ml/kg). For all minipigs, including the 4 female F1 animals, CSFs from the first sampling were used for analyses. CSF from one of the cloned *SORL1-het* animals (6402) was, however, excluded from the analyses due to contamination with blood during sampling. The brains were immediately and carefully removed from the scull and immersed in 4% phosphate-buffered formaldehyde for 2 weeks after which formaldehyde was replaced with phosphate-buffered saline and storage at 4°C.

### RT-PCR

Cerebellum and cortex samples were dissected from a *SORL1*^+/+^, a *SORL1*^+/−^ and a *SORL1*^−/−^ minipig, respectively, and RNA was extracted using the RNeasy Mini Kit (Qiagen, USA). 1 μg of total RNA was subsequently converted to cDNA using High Capacity RNA-to-cDNA Kit (#4387406, Applied Biosystems, USA) following manufacturer’s instructions. The resulting cDNA was used as template for amplification of porcine *SORL1*, using primer pairs located in the 5’end (exon 1-2) and in the 3’end (exon 46-47) of the transcript. GAPDH served as control for successful cDNA synthesis whereas samples without inclusion of reverse transcriptase (-RT) and water were used as negative controls. PCR was performed on a Veriti Thermal Cycler (Applied Biosystems) with Herculase II Fusion DNA Polymerase (#600675, Agilent) according to the following optimized conditions: 95°C for 1 min, 30 cycles of amplification (95°C for 20 sec, 65°C for 20 sec, 68°C for 1 min), and final extension at 68°C for 4 min. All primers used for RT-PCR are listed in **Supplemental Table S2**.

### Western blotting

Following dissection of cerebellum and cortex samples from *SORL1*^+/+^, *SORL1*^+/−^ and *SORL1^−/^* minipigs, tissues were homogenized and lysed. Separation of an equal amount of proteins by SDS-PAGE was performed using 4-12% NuPAGE Tris-Acetate gels (Invitrogen). Subsequently, proteins were transferred to nitrocellulose membrane (Amersham) in blotting buffer (Tris-base 250 mM, glycine 1.92 M) for 2.5 hs. Blocking of unspecific binding to the membrane was performed for 1h at room temperature in blocking buffer (TST buffer 20% (Tris-base 0.25 M, NaCl 2.5 M, 0.5% Tween-20, pH 9.0), 2% skimmed milk powder, and 2% Tween-20). The membrane was incubated overnight at 4°C with mouse anti-LR11 (1:500, #612633, BD Transduction Laboratories) for SORLA detection, and with mouse anti-actin (1:1000, #A5441, Sigma) as loading control. Next, the membrane was washed three times for 5 min in washing buffer (CaCl_2_ 0.2 mM, MgCl_2_ 0.1 mM, HEPES 1 mM, NaCl 14 mM, 0.2% skimmed milk powder, 0.05% Tween-20) and incubated with HRP-conjugated secondary antibody (1:1500, #P0260, Agilent) for 1h at RT. Proteins were finally detected with ECL detection reagent (GE Healthcare Amersham) and visualized with LAS-4000 (Fujifilm). For minipig CSF, equal volumes of CSF (10 uL) were loaded and Western blotting was performed as described above for proteins isolated from tissue homogenates.

### Mesoscale discovery

The Aβ standards (Aβ1-38, Aβ1-40 and Aβ-42) were diluted in Diluent 35 in a 4-fold serial dilution as instructed from the manufacturer (for MSD MULTI-SPOT Aβ Peptide Panel 1 V-Plex Assay). The precoated MSD plate was blocked in Diluent 35 for 1 hour at room temperature followed by washing in 1X PBS + 0,05% Tween-20. Detection antibody and standards/CSF samples were added to the wells, and incubated at room temperature for 2 hours. Lastly, the wells were washed and 2X Read Buffer T was added for immediate analysis. Electrochemiluminescent signals were obtained on a SECTOR® Imager (MSD) and the Aβ concentrations were calculated in Microsoft Excel by fitting the samples to a standard curve.

### Neurofilament light polypeptide assay

From the CSF that was collected from anesthetized minipigs at necropsy (10 wild-type animals and 6 *SORL1*-compromised minipigs (*SORL1*^+/−^, N=5 and *SORL1*^−/−^, N=1)), aliquots of 15 μL each were prepared and immediately frozen on dry ice prior to storage at −80°C until analysis for neurofilament light polypeptide (NF-L) levels. NF-L levels were quantified at a 1:50 dilution using the Protein Simple Platform ELLA according to the manufacturer’s instructions (LLOQ: 2.7 pg/ml, ULOQ: 10 290 pg/ml). Parallelism was observed for CSF dilutions between 1:10 and 1:100. For analysis, animals were grouped by genotype (*wt* versus *het*+*ko*), independent of age.

### Pathology

Neuropathological examination was performed blinded, initially macroscopically on slices of the formalin-fixed brains, and subsequently microscopically on formalin-fixed, paraffin-embedded and hematoxylin-stained brain sections from the frontal, temporal and occipital lobes, from the brain stem (mesencephalon, pons and myelencephalon) and cerebellum obtained from 2 F1 female minipigs (1 *het* and 1 *wt*) at 5 months of age, respectively.

### Immunohistochemistry

For the SORLA expression analysis in porcine neurons, paraffin embedded brain tissue was sectioned on a microtome (Leica RM2155) with 5 μm thickness followed by heating at 60°C for 1 hour. Sections of tissue were then deparaffinated in a xylene substitute (#109843, Neo-Clear, Merck) followed by dehydration in a series of ethanol (absolute to 70%) and rinsed in TBS (Tris buffered saline, pH 7.8) for 5 min. at room temperature. The DAKO Real™ EnVision™ System (#K4010, Dako) was used for the immunostaining. The sections were treated with an antigen retrieval solution (Target Retrieval pH 9, #S2367, DAKO) for 20 min. in a microwave and rinsed in TBS for 5 min at room temperature. The antigen retrieval was followed by incubation with peroxidase block for 5 min at room temperature and rinsed in TBS for 5 min. at room temperature. The sections were then incubated with a primary in-house antibody (SORLA rabbit pAb 5387) at 4°C over night and rinsed in TBS for 5 min at room temperature. The sections were incubated with secondary antibody (Peroxidase Labelled Polymer Anti-rabbit; #P0217, Agilent) for 30 min. at room temperature and rinsed in TBS for 5 min. at room temperature. For detection, a solution of Liquid DAB+ Chromogen in Substrate Buffer was used. The sections were incubated for 10 min. at room temperature, rinsed in TBS for 5 min at room temperature followed by staining with hematoxylin (#MHS16, Sigma Aldrich) for 3 min. at room temperature. The sections were then washed in deionized water (ddH_2_O), dehydrated in a series of ethanol (50% to absolute) and treated with a xylene substitute (#109843, Neo-Clear, Merck). Cover glass were mounted with Eukitt mounting medium (#03989, Sigma Aldrich) and images were acquired using a standard bright-field microscope (Zeiss Apotome) (**Fig. 1a**).

### [^18^F]-FDG and [^11^C]-PIB-PET imaging

Four 21-month old female minipigs (*wt*, N=2; *het*, N=2) underwent PET neuroimaging with [^18^F]-FDG and [^11^C]-PIB at the Department of Nuclear Medicine and PET at Aarhus University Hospital. The 16 hours fasted minipigs were premedicated with an intramuscular injection of 1.3 mg/kg midazolam and 6.3 mg/kg s-ketamine. Anaesthesia was then induced through an ear vein catheter with 1.3 mg/kg midazolam and 3.1 mg/kg s-ketamine. After intubation, the anaesthesia was maintained with 2.0-2.1 % isoflurane. The pigs were mechanically ventilated 15 times per minute with approximately 8 ml/kg of a 1: 2.2 oxygen-medical air mixture. The minipigs were placed prone with their heads in the center of the field of view of a Siemens Biograph Truepoint PET/CT system. A CT scan for attenuation correction of PET data was acquired. For [^18^F]-FDG, a dose of 5MBq/kg of was then injected via the ear vein and 45 minutes later, a 30-minute PET scan was acquired. The injected dose in the 4 pigs ranged from 170-216MBq depending on bodyweight. Similarly, on a separate day, 138-204MBq of [^11^C]-PIB was injected into the ear vein and 90 min dynamic imaging began at tracer injection. Both tracers were administered in 20 mls of saline and then flushed with 10 mls of saline. After [^11^C]-PIB injection, the catheter was removed to avoid scanner artifacts. PET images were reconstructed using the iterative TrueX algorithm (Siemens), and CT and PET data were fused by the system. PET scans were then co-registered to an average MRI of the minipig brain using PMOD version 4.0 and regional standard uptake values were obtained and normalized to whole brain activity.

### Magnetic resonance imaging

Four female 22-month old minipigs (*wt*, N=2; *het*, N=2) and four 27-month old male minipigs (*wt*, N=2; *het*, N=2) were subjected to Magnetic Resonance Imaging (MRI). The pigs were prepared in the same ways as prior to PET imaging, but instead of isoflurane, the anaesthesia was maintained with intravenous propofol (approximately 8 mg/kg/hour) infusion. MRI was carried out on a 3T scanner (MR750, GE Healthcare) in one session. We obtained structural proton images and DTI for assessment of volumes and diffusivity using a flexible 16-channel coil (GE Healthcare). Anatomy and volumes were evaluated using T_1_-weighted (TR/TE = 14/500 ms, 0.8 × 0.8 × 1.4 mm^3^) and T_2_-weighted (TR/TE/TI = 6300/103/1750 ms, 0.8 × 0.8 × 1.4 mm^3^) fluid attenuation inversion recovery (FLAIR) 3D imaging. Further, we performed DTI (TR/TE = 8000/65 ms, 30 directions, b = 1000 s/mm^2^, 2.1 × 1.9 × 4 mm^3^), which was processed and analyzed using FSL (Analysis Group, FMRIB, Oxford) (51).

### Statistics

All statistical tests were done using the program Prism GraphPad version 6.0. For comparison between two different datasets a two-tailed unpaired Student’s *t*-test was used. *P* values below 0.05 were considered significant. Error bars represent standard errors of the mean.

## Acknowledgement

We would like to thank Aage Kristian Olsen Alstrup, Dept. of Clinical Medicine, Aarhus University/Dept. of Nuclear Medicine & PET, Aarhus University Hospital, Denmark, and Kjeld Dahl Winther, SEGES, Denmark, for excellent veterinary assistance during PET/MRI scannings and CSF sampling, respectively. Also, we are grateful to Bert Vogelstein and Kenneth W. Kinzler, Johns Hopkins University, Baltimore, Maryland, for kindly providing the pNeDaKO plasmid vector. We would also like to thank Trine S. Petersen, Lisa Maria Røge, Dorte Qualmann, Anette M. Pedersen, Janne Adamsen, Klaus Villemoes, Michelle Sørensen, Sandra Bonnesen, and Benedicte Vestergaard for skilled technical assistance, and Anne Mette V. Toft, Mette Bak and Martin A. Fredsted for skillful assistance with animal handling and transport to scanning facilities. This study was carried out in accordance to the ARRIVE guidelines and was supported by grants to C.B.S. from the Lundbeck Foundation (R100-A9209), The Danish Heart Association (16-R107-A6813-22997), Ellegaard Göttingen Minipigs Research Foundation, and Ellegaard Göttingen Minipigs A/S.

## Disclosures

LB and MD are employees of AbbVie and own AbbVie stock. AbbVie participated in the design, study conduct, and financial support for this research as well as in the interpretation of data, review, and approval of the publication.

Ellegaard Göttingen Minipigs A/S is having the commercialization rights to the genetically altered Göttingen Minipigs *SORL1* KO model.

OMA and SAS has commercial interests in Retromer Therapeutics, but this company was not involved in any aspects of the current study.

## SUPPLEMENTAL INFORMATION

### List of Supplementary information

Supplementary Figures S1-S6

Supplementary Table S1-S2

**Table S1. Animals included in the study**

**Table S2. Primer sequences**

**Figure S1. Molecular genetic validation of the *SORL1* Göttingen Minipigs**

a. Southern blot
b. Genotyping

**Figure S2. Alignment of human and pig SORLA protein sequences**

**Figure S3. Histopathology of young *SORL1-het* and *SORL1-wt* Göttingen Minipigs**

Histological examination of hematoxylin-stained frontal cortex from 5-month old *wt* (6478) and *het* (6473) minipigs showing no detectable difference in neuronal layering and numbers.

**Figure S4. Alignment of human and porcine APP protein sequences**

**Figure S5. Alignment of human and porcine tau protein sequences**

**Figure S6. Normal neuronal integrity in *SORL1-het* and *SORL1-wt* Göttingen Minipigs**

Biochemical assessment of neuronal degradation as measured by neurofilament light chain (NF-L) in collected minipig CSFs (*wt*, N=10; *het*, N=5; *ko*, N=1). The group of *het* minipigs were depicted including data obtained from the *ko* pig 6304 shown in red.

