## Supplemental information for "*In vivo* evidence that *SORL1*, encoding the endosomal recycling receptor SORLA, can function as a causal gene in Alzheimer’s Disease"

**Supplemental Table 1. Overview of pigs included in the study.**

| Pig (ID no) | Sex | Genotype | Age at CSF sampling (mo) | Age at FDG-PET scan (mo) | Age at PIB-PET scan (mo) | Age at MRI scan (mo) | Age at euthanization (mo) |
|---|---|---|---|---|---|---|---|
| **Cloned animals (F0)** | | | | | | | |
| 6304 | Female | KO | 33 | - | - | - | 33 |
| 6401 | Female | HET | 35 | - | - | - | 35 |
| 6402 | Female | HET | 36* | - | - | - | 36 |
| **F1 offspring** | | | | | | | |
| 6469 | Female | HET | 24 | 21 | 21 | 22 | 29 |
| 6470 | Female | HET | 24 | 21 | 21 | 22 | 29 |
| 6471 | Male | WT | 18 | - | - | - | 18 |
| 6472 | Male | HET | 30 | - | - | 27 | 30 |
| 6473 | Female | HET | 5 | - | - | - | 5 |
| 6474 | Male | HET | 30 | - | - | 27 | 30 |
| 6475 | Female | WT | 24 | 21 | 21 | 22 | 29 |
| 6476 | Male | HET | 18 | - | - | - | 18 |
| 6477 | Female | WT | 24 | 21 | 21 | 22 | 29 |
| 6478 | Female | WT | 5 | - | - | - | 5 |
| **WT control animals** | | | | | | | |
| 334011 | Female | WT | 36 | - | - | - | 36 |
| 230251 | Female | WT | 38 | - | - | - | 38 |
| 339671 | Male | WT | 17 | - | - | - | 17 |
| 339704 | Male | WT | 17 | - | - | - | 17 |
| 338496 | Male | WT | 29 | - | - | 27 | 29 |
| 338593 | Male | WT | 30 | - | - | 27 | 29 |
| **WT breeding boars** | | | | | | | |
| 229751 | Male | WT | 23 | - | - | - | 23 |
| 332725 | Male | WT | 22 | - | - | - | 22 |

Pigs included in the study listing age in whole months at individual procedures. A dash indicate that the procedure was not performed for the animal in question.

* CSF from animal 6402 was contaminated with blood and not used for any analysis

**Supplemental Table S2. Primer sequences**

| | Forward primer (5´- 3´) | Reverse primer (5´- 3´) |
|---|---|---|
| **rAAV/*SORL1* KO vector** | | |
| LHA* | atacatacgcggccgcCCTCAAAAACCAGGGTGTGAGTCCAGAGC | *gctccagcttttgttccctttag*CCCTGGCTGGCGCTCCTCCTTGTCCG |
| RHA* | *cgccctatagtgagtcgtattac*GCATCCATCCTTGGCTGTCGCTTCCCAGG | atacatacgcggccgcCCAAGGGTTGAACATTTAACTCTGTGTATTTCC |
| 3-Fusion* | atacatacgcggccgcCCTCAAAAACCAGGGTGTGAGTCCAGAGC | atacatacgcggccgcCCAAGGGTTGAACATTTAACTCTGTGTATTTCC |
| **CRISPR sgRNA vectors** | | |
| sgRNA1** | accgGCGACACGGAGCAGCAGGA | aaacTCCTGCTGCTCCGTGTCGC |
| sgRNA2** | accgATGGCGCTGCTGCCGCCCG | aaacCGGGCGGCAGCAGCGCCAT |
| **C-check vector** | | |
| sgRNA1-sgRNA2 target site insert*** | gtcggat(GGCGACACGGAGCAGCAGGAGGGGATGGCGCTGCTGCCGCCCGGGG)aggt | cggtacct(CCCCGGGCGGCAGCAGCGCCATCCCCTCCTGCTGCTCCGTGTCGCC)atc |
| **PCR screening of donor cells** | | |
| 5´ *SORL1* KO PCR (F3+R3) | CTCTAGAAGTAGTCTCTCCTTCAGTCCTG | GCGCATGCTCCAGACTGCCTTGG |
| 3´ *SORL1* KO PCR (F4+R4) | GGTACCCAATTCGCCCTATAGTGAGTC | AATGACACAATAGGCTAAGATGG |
| **Southern blot probes** | | |
| *SORL1* probe | CTGAGCTCCCCAAAGTTAGAAAGTG | GCCTCTCCAGTTAACAGACCTTCC |
| neo$^r$ probe | GAAGCCCGGCATTCTGCACGC | CAGAAGCCATAGAGCCCACCGCA |
| **Off target analysis** | | |
| Chr. 2 (*JUNB*) | AGCGGTGGCGGCAGCTAC | GTACGAGCTCCCGGTACCGAC |
| Chr. 2 (ARHGAP26) | AGCGCCAGGAGGCCCATG | CAAGGATGCGCTCCGTTCG |
| Chr. 5 (*XRCC6*) | CCGTGCCACTCACCATTCACC | CAGTTCACTCGTGTGACCTGGAGC |
| Chr. 6 (*GSE1*) | CTGAATCAGCACATGTCTGGCC | GGACCTCGCCGTGAGCAG |
| Chr. 8 (*PCDH7*) | GAGTGGGATACGACCGATCTTGC | GCGAACAGCGCGACTCCTATG |
| Chr. 9 (*HEPACAM*) | CTCAGGCGCTGATCTCCACAG | GTGCAGACTTCCAGGAGAGGTCC |
| Chr. 14 (*TXNRD2*) | CCTGGAGGTTCACAGCAAGGTC | GGTACCCTGAGTGAGCTTTGCTC |
| Chr. 15 (*TWIST2*) | AGGCATGACCAGGTCATTCAGG | TCACGGAGGGAGCTCGGC |
| **Plasmid integration analysis** | | |
| sgRNA1 plasmid | GGACATAAGCCTGTTCGGTTCG | ACGCCACGGAATGATGTCG |
| hCas9 plasmid | ATGGGCGGTAGGCGTGTAC | GATCTCCTGCAGGTAGCAGATC |
| **Genotyping of piglets** | | |
| 5´ *SORL1* KO PCR | CTCTAGAAGTAGTCTCTCCTTCAGTCCTG | GCGCCTACCGGTGGATGTGG |
| 3´ *SORL1* KO PCR | GGTACCCAATTCGCCCTATAGTGAGTC | AATGACACAATAGGCTAAGATGG |
| *SORL1* WT PCR | CTCTAGAAGTAGTCTCTCCTTCAGTCCTG | CTTCGCGCACTTTCTCCGCTG |
| **RT-PCR** | | |
| 5´ *SORL1* RT-PCR (exon 1-2, F1+R2) | CGGACGAGAAGCCGCTCCG | CAAGGCCACGATGACGTTGC |
| 5´ *SORL1* RT-PCR (exon 1-3, F1+R1) | CGGACGAGAAGCCGCTCCG | GCGATCACGGCTTCACTGCTG |
| 3´ *SORL1* RT-PCR (exon 46-47) | GCCATGAACATCACAGCGTACC | GCTGACACATCCCTACCAGACC |
| *GAPDH* RT-PCR | GACTCATGACCACGGTCCATG | GTCAGATCCACAACCGACACG |

*Underlined sequence: *Not*I restriction site; sequence in italics: linker for 3-fusion PCR
**Lower case: overhang for BsaI cloning; upper case: porcine *SORL1* target sequence
***Nucleotides in brackets denote the overlapping target site sequences recognized by sgRNA1 and sgRNA2. Nucleotides relevant for cloning are shown in lower case.

Andersen
Supplemental Figure S1

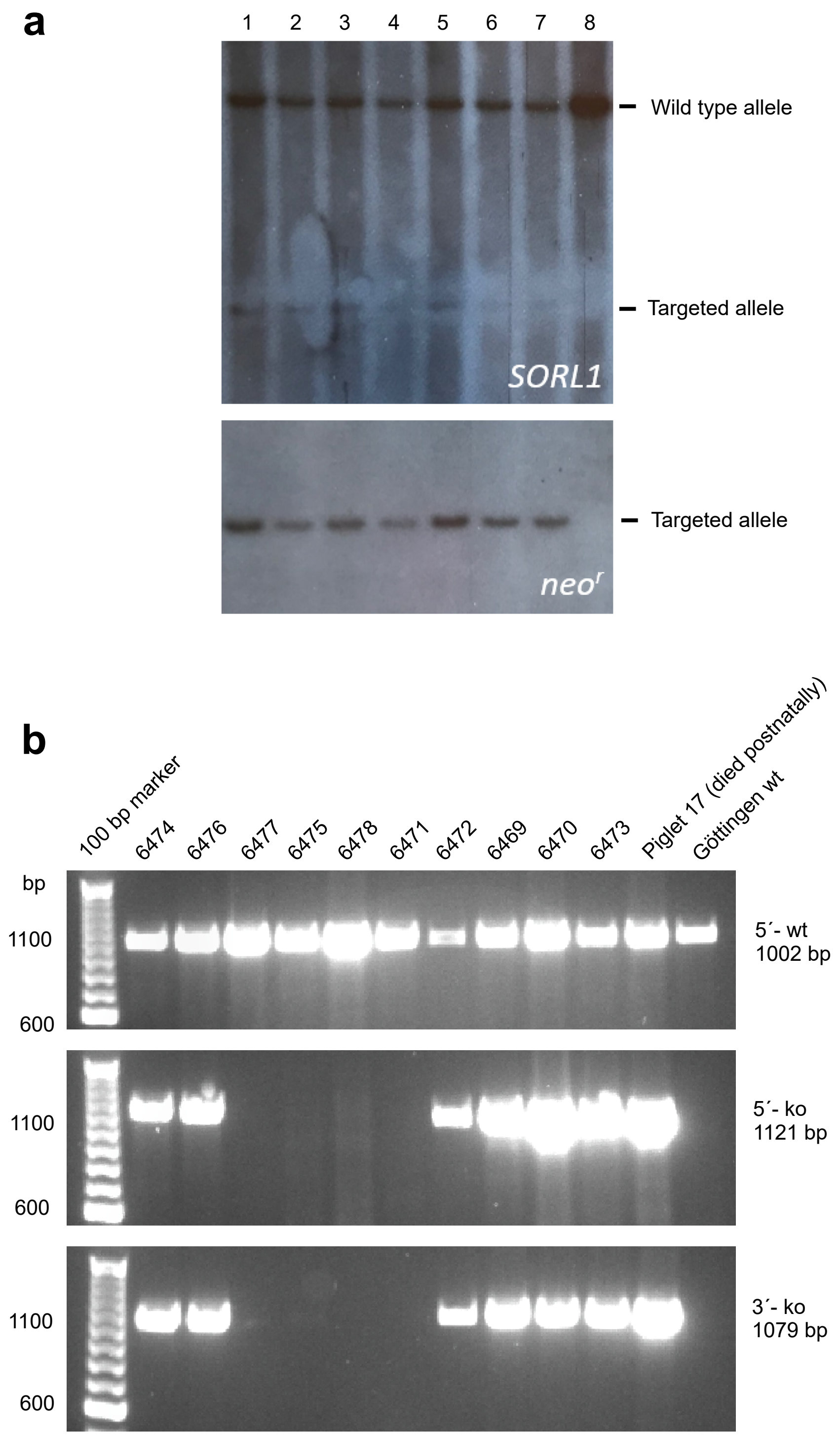

**Suppl Figure S2**
**SORLA identity 92.2% (2043/2216)**

```
                         10        20        30        40        50        60
  Pig      MATRSSRRESRLPFLFILMALLPPGAVGEVWPYTLPGGRAPGPQDRGFLVVRGDPRELRL
Human      MATRSSRRESRLPFLFTLVALLPPGALCEVWTQRLHGGSAPLPQDRGFLVVQGDPRELRL

                         70        80        90       100       110       120
  Pig      GAHGAPRG---ADEKPLRRKRSAALQPEPIQVYGQVSLNDSHNQMVVHWAGEKSNVIVAL
Human      WARGDARGASRADEKPLRRKRSAALQPEPIKVYGQVSLNDSHNQMVVHWAGEKSNVIVAL

                        130       140       150       160       170       180
  Pig      ARDSLSLARPRSSDVYVSYDYGRSFKRISEKLNFGTGNSSEAVIAQFYHSPADNKRYIFA
Human      ARDSLALARPKSSDVYVSYDYGKSFKKISDKLNFGLGNRSEAVIAQFYHSPADNKRYIFA

                        190       200       210       220       230       240
  Pig      DAYAQYLWTTFDFCNTIQGFSIPFRAADLLLHSKASNLLLGFDRSHPNKQLWKSDDFGQT
Human      DAYAQYLWITFDFCNTLQGFSIPFRAADLLLHSKASNLLLGFDRSHPNKQLWKSDDFGQT

                        250       260       270       280       290       300
  Pig      WIMIQEHVKSFSWGVDPYDKPNTIYVERHEPSGYSTVFRSTDFFQSRENLEVILEEVRDF
Human      WIMIQEHVKSFSWGIDPYDKPNTIYIERHEPSGYSTVFRSTDFFQSRENQEVILEEVRDF

                        310       320       330       340       350       360
  Pig      QLRDKYMFATKVVQHLFGSQQPSSVQLWVSFGRKPMRAAQFVTRHPINEYYIADASEDQV
Human      QLRDKYMFATKVV-HLLGSEQQSSVQLWVSFGRKPMRAAQFVTRHPINEYYIADASEDQV

                        370       380       390       400       410       420
  Pig      FVCVSHSNNRTNLYISEAEGLKFSLSLENVLYYSPGGAGSDTLVRYFANEPFADFHRVEG
Human      FVCVSHSNNRTNLYISEAEGLKFSLSLENVLYYSPGGAGSDTLVRYFANEPFADFHRVEG

                        430       440       450       460       470       480
  Pig      LQGVYIATLINGSMNEENMRSVITFDKGGTWEFLQAPAFTEYGEKINCELSQGCSLHLAQ
Human      LQGVYIATLINGSMNEENMRSVITFDKGGTWEFLQAPAFTGYGEKINCELSQGCSLHLAQ

                        490       500       510       520       530       540
  Pig      RLSQLLSLQLRRTPILSKESAPGLIIATGSVGKNLASKTNVYVSSSAGARWREALPGPHY
Human      RLSQLLNLQLRRMPILSKESAPGLIIATGSVGKNLASKTNVYISSSAGARWREALPGPHY

                        550       560       570       580       590       600
  Pig      YTWGDHGGIIVAIAQGMETNELKYSTNEGETWKTFVFSEKPMFVYGLLTEPGEKSTVFTI
Human      YTWGDHGGIITAIAQGMETNELKYSTNEGETWKTFIFSEKPVFVYGLLTEPGEKSTVFTI

                        610       620       630       640       650       660
  Pig      FGSNKENIHSWLILQVNTTDALGVPCTENDYKLWSPSDERGNECLLGHKTVFKRRTPHAT
Human      FGSNKENVHSWLILQVNATDALGVPCTENDYKLWSPSDERGNECLLGHKTVFKRRTPHAT

                        670       680       690       700       710       720
  Pig      CFNGEDFDRPVVVSNCSCTREDYECDFGFKMSEDLFLEVCVPDPEFSGKSFSPPVPCPVG
Human      CFNGEDFDRPVVVSNCSCTREDYECDFGFKMSEDLSLEVCVPDPEFSGKSYSPPVPCPVG

                        730       740       750       760       770       780
  Pig      STYRRTRGYRKISGDTCSGGDVEVRLEGELVPCPLAEENEFILYAMRKSIHRYDLASGTT
Human      STYRRTRGYRKISGDTCSGGDVEARLEGELVPCPLAEENEFILYAVRKSIYRYDLASGAT

                        790       800       810       820       830       840
  Pig      EQLPLTGLRAAVALDFDYEHNCLYWSDLALDIIQRLCLNGNTGQEVIINSGLETVEALAF
Human      EQLPLTGLRAAVALDFDYEHNCLYWSDLALDVIQRLCLNGSTGQEVIINSGLETVEALAF

                        850       860       870       880       890       900
  Pig      EPLSQLLYWVDSGFKKIEVGNPDGDFRLTIVNSSVLDRPRALVLVPQDGVMFWTDWGDLR
Human      EPLSQLLYWVDAGFKKIEVANPDGDFRLTIVNSSVLDRPRALVLVPQEGVMFWTDWGDLK

                        910       920       930       940       950       960
  Pig      PGIYRSNMDGSAAYRLVSEDVKWPNGIAVDAQWVYWTDAYLDCIERVTFSGQQRSIILDN
Human      PGIYRSNMDGSAAYHLVSEDVKWPNGISVDDQWIYWTDAYLECIERITFSGQQRSVILDN

                        970       980       990      1000      1010      1020
  Pig      LPHPYAIAVFKNEIYWDDWSELSIFRASKHSKSDMAILASQLTGPMDLKIFYRGKTTGSN
Human      LPHPYAIAVFKNEIYWDDWSQLSIFRASKYSGSQMEILANQLTGLMDMKIFYKGKNTGSN

                       1030      1040      1050      1060      1070      1080
  Pig      ACVSRPCSLLCLPKANNTRTCRCPDGVSGSVLPSGDLMCECPQGYQLKNHTCVKEENTCL
Human      ACVPRPCSLLCLPKANNSRSCRCPEDVSSSVLPSGDLMCDCPQGYQLKNNTCVKEENTCL

                       1090      1000      1110      1120      1130      1140
  Pig      RNQYRCSNGNCINSIWWCDFDNDCGDMSDERNCPTTVCDLDTQFRCQESGTCIPLSYKCD
Human      RNQYRCSNGNCINSIWWCDFDNDCGDMSDERNCPTTICDLDTQFRCQESGTCIPLSYKCD

                       1150      1160      1170      1180      1190      1200
  Pig      LEDDCGDNSDESHCELHQCRSNEYSCSSGMCIRSSWVCDGDNDCRDWSDEANCTAIYHTC
Human      LEDDCGDNSDESHCEMHQCRSDEYNCSSGMCIRSSWVCDGDNDCRDWSDEANCTAIYHTC

                       1210      1220      1230      1240      1250      1260
  Pig      EASNFQCHNGHCIPQRWACDGDMDCQDGSDEDPVTCEKKCNGFRCPNGTCIPSSKHCDGL
Human      EASNFQCRNGHCIPQRWACDGDTDCQDGSDEDPVNCEKKCNGFRCPNGTCIPSSKHCDGL
```

```
             1270      1280      1290      1300      1310      1320
Pig    HDCGDGSDEQHCEPLCTRFMDFVCKNRQQCLFQSMVCDGIVQCRDGSDEDAAFAGCSHDP
Human  RDCSDGSDEQHCEPLCTHFMDFVCKNRQQCLFHSMVCDGIIQCRDGSDEDAAFAGCSQDP

             1330      1340      1350      1360      1370      1380
Pig    EFHKVCDELSFQCQNGVCISLIWKCDGMDDCGDDSDEANCENPTEAPNCSRYFQFQCENG
Human  EFHKVCDEFGFQCQNGVCISLIWKCDGMDDCGDYSDEANCENPTEAPNCSRYFQFRCENG

             1390      1400      1410      1420      1430      1440
Pig    HCIPNRWKCDRENDCGDWSDEKDCGDLHILPSPTPGPSTCLPNYYRCSSGACVMDSWVCD
Human  HCIPNRWKCDRENDCGDWSDEKDCGDSHILPFSTPGPSTCLPNYYRCSSGTCVMDTWVCD

             1450      1460      1470      1480      1490      1500
Pig    GYRDCADGSDEEACPSPANVTAASTPTQLGRCDRFEFECRQPKKCIPNWRRCDGHQDCQD
Human  GYRDCADGSDEEACPLLANVTAASTPTQLGRCDRFEFECHQPKTCIPNWKRCDGHQDCQD

             1510      1520      1530      1540      1550      1560
Pig    GQDEANCPTRSSLTCTSWEFKCEDGETCIVLSERCDGFLDCSDESDERNCSEELNVYKIQ
Human  GRDEANCPTHSTLTCMSREFQCEDGEACIVLSERCDGFLDCSDESDEKACSDELTVYKVQ

             1570      1580      1590      1600      1610      1620
Pig    NLQWTADFSGDITLTWLKPKKMPSASCVYNVYYRVVGESMWKILETHSNKTSTVLKVLKP
Human  NLQWTADFSGDVTLTWMRPKKMPSASCVYNVYYRVVGESIWKTLETHSNKTNTVLKVLKP

             1630      1640      1650      1660      1670      1680
Pig    DTTYQVKVQVQCLSKVHSTNDFVTLRTPEGLPDAPQNLQLSLHREVEGVIVGHWTPPIHT
Human  DTTYQVKVQVQCLSKAHNTNDFVTLRTPEGLPDAPRNLQLSLPREAEGVIVGHWAPPIHT

             1690      1700      1710      1720      1730      1740
Pig    HGLIREYIVEYSRSGSKMWASQRSAGNSTEIRNLLLNAPYTVRVAAVTSRGIGNWSDSKS
Human  HGLIREYIVEYSRSGSKMWASQRAASNFTEIKNLLVNTLYTVRVAAVTSRGIGNWSDSKS

             1750      1760      1770      1780      1790      1800
Pig    ITTTNKGKVIPQPDIHIDSYDENSLSFTLSMDSDIKVNGYVVNLFWAFDTHAQEQRTLNF
Human  ITT-IKGKVIPPPDIHIDSYGENYLSFTLTMESDIKVNGYVVNLFWAFDTHKQERRTLNF

             1810      1820      1830      1840      1850      1860
Pig    QGSMLSHRVGNLTAHTPYEISAWAKTDLGDSPLAFEHVTTRGVRPPAPSLKAKAINQTAV
Human  RGSILSHKVGNLTAHTSYEISAWAKTDLGDSPLAFEHVMTRGVRPPAPSLKAKAINQTAV

             1870      1880      1890      1900      1910      1920
Pig    ECTWTGPRNVVYGIFYATSFLDLYRNPKSVTTSQHNKTVLVSRDEQYLFLVRVVVPYQGP
Human  ECTWTGPRNVVYGIFYATSFLDLYRNPKSLTTSLHNKTVIVSKDEQYLFLVRVVVPYQGP

             1930      1940      1950      1960      1970      1980
Pig    SSDYVVVRMIPDSRLPPRHLHVVHTGKTSAVIKWESPYDSPDQDLLYAIAVKDLIRKSDR
Human  SSDYVVVKMIPDSRLPPRHLHVVHTGKTSVVIKWESPYDSPDQDLLYAIAVKDLIRKTDR

             1990      2000      2010      2020      2030      2040
Pig    SYKVKSRNSTVEYTLNKLEPGGKYHVIVQLGNMSKDSSIKITTVSLSAPDALKIITENDH
Human  SYKVKSRNSTVEYTLNKLEPGGKYHIIVQLGNMSKDSSIKITTVSLSAPDALKIITENDH

             2050      2060      2070      2080      2090      2100
Pig    VLLFWKSLALKEKHFNESRGYEIHMFDSAMNITAYLGNTTDNFFKISNLKLGHNYTFTVQ
Human  VLLFWKSLALKEKHFNESRGYEIHMFDSAMNITAYLGNTTDNFFKISNLKMGHNYTFTVQ

             2110      2120      2130      2140      2150      2160
Pig    AQCLFGSQICGEPAVLLYDELGSGRDVSAIQATRSTDVAAVVVPILFLILLSLGIGFAIL
Human  ARCLFGNQICGEPAILLYDELGSGADASATQAARSTDVAAVVVPILFLILLSLGVGFAIL

             2170      2180      2190      2200      2210
Pig    YTKHRRLQNSFTAFANSHYSSRLGSAIFSSGDDLGEDDEDAPMITGFSDDVPMVIA*
Human  YTKHRRLQSSFTAFANSHYSSRLGSAIFSSGDDLGEDDEDAPMITGFSDDVPMVIA*
```

Andersen
Supplemental Figure S3

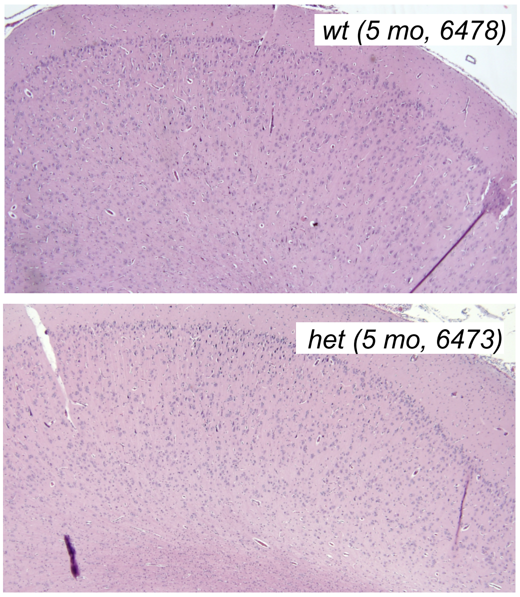

**Suppl Figure S4**
**APP identity 97.8% (753/770)**

```
                 10        20        30        40        50        60
Pig      MLPGLALVLLAAWTARALEVPTDGNAGLLAEPQVAMFCGKLNMHMNVQNGKWESDPSGTK
Human    MLPGLALLLLAAWTARALEVPTDGNAGLLAEPQIAMFCGRLNMHMNVQNGKWDSDPSGTK

                 70        80        90       100       110       120
Pig      TCIGTKEGILQYCQEVYPELQITNVVEANQPVTIQNWCKRSRKQCKTHTHIVIPYRCLVG
Human    TCIDTKEGILQYCQEVYPELQITNVVEANQPVTIQNWCKRGRKQCKTHPHFVIPYRCLVG

                130       140       150       160       170       180
Pig      EFVSDALLVPDKCKFLHQERMDVCETHLHWHTVAKETCSEKSTNLHDYGMLLPCGIDKFR
Human    EFVSDALLVPDKCKFLHQERMDVCETHLHWHTVAKETCSEKSTNLHDYGMLLPCGIDKFR

                190       200       210       220       230       240
Pig      GVEFVCCPLAEESDNIDSADAEEDDSDVWWGGADTDYADGSEDKVVEVAEEEEVADVEEE
Human    GVEFVCCPLAEESDNVDSADAEEDDSDVWWGGADTDYADGSEDKVVEVAEEEEVAEVEEE

                250       260       270       280       290       300
Pig      EAEDDEDDEDGDEVEEEAEEPYEEATERTTSIATTTTTTTESVEEVVREVCSEQAETGPC
Human    EADDDEDDEDGDEVEEEAEEPYEEATERTTSIATTTTTTTESVEEVVREVCSEQAETGPC

                310       320       330       340       350       360
Pig      RAMISRWYFDVTEGKCAPFFYGGCGGNRNNFDTEEYCMAVCGSVMSQSLLKTTQEHLPQD
Human    RAMISRWYFDVTEGKCAPFFYGGCGGNRNNFDTEEYCMAVCGSAMSQSLLKTTQEPLARD

                370       380       390       400       410       420
Pig      PVKLPTTAASTPDAVDKYLETPGDENEHAHFQKAKERLEAKHRERMSQVMREWEEAERQA
Human    PVKLPTTAASTPDAVDKYLETPGDENEHAHFQKAKERLEAKHRERMSQVMREWEEAERQA

                430       440       450       460       470       480
Pig      KNLPKADKKAVIQHFQEKVESLEQEAANERQQLVETHMARVEAMLNDRRRLALENYITAL
Human    KNLPKADKKAVIQHFQEKVESLEQEAANERQQLVETHMARVEAMLNDRRRLALENYITAL

                490       500       510       520       530       540
Pig      QAVPPRPRHVFNMLKKYVRAEQKDRQHTLKHFEHVRMVDPKKAAQIRSQVMTHLRVIYER
Human    QAVPPRPRHVFNMLKKYVRAEQKDRQHTLKHFEHVRMVDPKKAAQIRSQVMTHLRVIYER

                550       560       570       580       590       600
Pig      MNQSLSLLYNVPAVAEEIQDEVDELLQKEQNYSDDVLANMISEPRISYGNDALMPSLTET
Human    MNQSLSLLYNVPAVAEEIQDEVDELLQKEQNYSDDVLANMISEPRISYGNDALMPSLTET

                610       620       630       640       650       660
Pig      KTTVELLPVNGEFSLDDLQPWHPFGVDSVPANTENEVEPVDARPAADRGLTTRPGSGLTN
Human    KTTVELLPVNGEFSLDDLQPWHSFGADSVPANTENEVEPVDARPAADRGLTTRPGSGLTN

                670       680       690       700       710       720
Pig      IKTEEISEVKMDAEFRHDSGYEVHHQKLVFFAEDVGSNKGAIIGLMVGGVVIATVIVITL
Human    IKTEEISEVKMDAEFRHDSGYEVHHQKLVFFAEDVGSNKGAIIGLMVGGVVIATVIVITL

                730       740       750       760       770
Pig      VMLKKKQYTSIHHGVVEVDAAVTPEERHLSKMQQNGYENPTYKFFEQMQN
Human    VMLKKKQYTSIHHGVVEVDAAVTPEERHLSKMQQNGYENPTYKFFEQMQN
```

**Suppl Figure S5**
**Tau identity 89.8%**

```
                                    10        20        30        40        50        60
  Pig            MAEPRQEFTVMEDHA----------QGDYTLLQDHEGDVDHSLKESPLQTPADDGSEEPG
Human            MAEPRQEFEVMEDHAGTYGLGDRKDQGGYTMHQDQEGDTDAGLKESPLQTPTEDGSEEPG

                                    70        80        90       100       110       120
  Pig            SETSDAKSTPTAEGDSWWFPDVTAPLVDEGAPGEQAAAQPPTEIPEGTTAEEAGIGDTPN
Human            SETSDAKSTPTAE-------DVTAPLVDEGAPGKQAAAQPHTEIPEGTTAEEAGIGDTPS

                                   130       140       150       160       170       180
  Pig            LEDQAAGHVTQARMVSKGKDGTGPDDKKTKGADGKPGSKIATPRGAAPPGQKGPANATRI
Human            LEDEAAGHVTQARMVSKSKDGTGSDDKKAKGADGK--TKIATPRGAAPPGQKGQANATRI

                                   190       200       210       220       230       240
  Pig            PAKTTPSPKTPPGSGESGKSGDRSGYSSPGSPGTPGSRSRTPSLPTPPTREPKKVAVVRT
Human            PAKTPPAPKTPPSSGEPPKSGDRSGYSSPGSPGTPGSRSRTPSLPTPPTREPKKVAVVRT

                                   250       260       270       280       290       300
  Pig            PPKSPSAAKSRLQTAPGPMPDLKNVRSKIGSTENLKHQPGGGKVQIINKKLDLSNVQSKC
Human            PPKSPSSAKSRLQTAPVPMPDLKNVKSKIGSTENLKHQPGGGKVQIINKKLDLSNVQSKC

                                   310       320       330       340       350       360
  Pig            GSKDNIKHVPGGGSVQIVYKPVDLSKVTSKCGSLGNIHHKPGGGQVEVKSEKLDFKDRVQ
Human            GSKDNIKHVPGGGSVQIVYKPVDLSKVTSKCGSLGNIHHKPGGGQVEVKSEKLDFKDRVQ

                                   370       380       390       400       410       420
  Pig            SKIGSLDNITHVPGGGNKKIETHKLTFRENAKAKTDHGAEIVYKSPVVSGDTSPRHLSNV
Human            SKIGSLDNITHVPGGGNKKIETHKLTFRENAKAKTDHGAEIVYKSPVVSGDTSPRHLSNV

                                   430       440       450
  Pig            SSTGSIDMVDSPQLATLADEVSASLAKQGL
Human            SSTGSIDMVDSPQLATLADEVSASLAKQGL
```

HT7
5E2

Thr-181 and Thr-217 are marked by green in the human Tau sequence

Andersen
Supplemental Figure S6

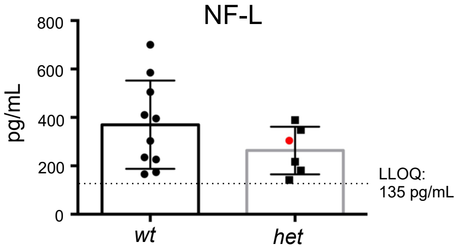
